# Hyperspectral Counting of Multiplexed Nanoparticle Emitters in Single Cells and Organelles

**DOI:** 10.1101/2021.11.24.469882

**Authors:** Prakrit V. Jena, Mitchell Gravely, Christian C. Cupo, Mohammad M. Safaee, Daniel Roxbury, Daniel A. Heller

## Abstract

Nanomaterials are the subject of a range of biomedical, commercial, and environmental investigations involving measurements in living cells and tissues. Accurate quantification of nanomaterials, at the tissue, cell, and organelle levels, is often difficult, however, in part due to their inhomogeneity. Here, we propose a method that uses the diverse optical properties of a nanomaterial preparation in order to improve quantification at the single-cell and organelle level. We developed ‘hyperspectral counting’, which employs diffraction-limited imaging *via* hyperspectral microscopy of a diverse set of nanomaterial emitters, to estimate nanomaterial counts in live cells and sub-cellular structures. A mathematical model was developed, and Monte Carlo simulations were employed, to improve the accuracy of these estimates, enabling quantification with single-cell and single-endosome resolution. We applied this nanometrology technique to identify an upper-limit of the rate of uptake into cells - approximately 3,000 particles endocytosed within 30 minutes. In contrast, conventional ROI counting results in a 230% undercount. The method identified significant heterogeneity and a broad non-Gaussian distribution of carbon nanotube uptake within cells. For example, while a particular cell contained an average of 1 nanotube per endosome, the heterogenous distribution resulted in over 7 nanotubes localizing within some endosomes, substantially changing the accounting of subcellular nanoparticle concentration distributions. This work presents a method to quantify cellular and subcellular concentrations of a heterogeneous carbon nanotube reference material, with implications for nanotoxicology, drug/gene delivery, and nanosensor fields.

---

In the fields of nanomedicine, nanotoxicology, and the environmental impact of nanotechnologies, characterizing the tissue and cellular uptake is of fundamental importance.^1–3^ The mechanism of uptake and subcellular localization of nanomaterials are typically predicted by their physicochemical properties,^4^ providing tunable parameters to control these processes. The delivered dose, *i.e.*, the quantity of internalized particles, is particularly relevant to nanotoxicity evaluations due to potential dose-dependent adverse effects.^5^ As a result, quantitative analyses are required for nanotoxicological conclusions to be drawn,^6^ yet fundamental advances in nanotoxicology have been hindered by a lack of standardization.^6–8^ The complex interactions which occur in biological settings create additional variables that can directly affect material uptake, including protein corona formation,^2^ which can be modulated by even minor experimental procedures.^9^ Minimum information reporting in bio-nano experimental literature (MIRIBEL) has therefore been suggested as a ‘minimum information standard’ to advance the principles of reusability, quantification, practicality and quality in material and biological characterization and experimental protocol details.^10^ Because analytical techniques rely on the intrinsic properties of nanomaterials, appropriate methodologies will inherently vary for specific materials,^8^ and thus a single approach cannot be used. Instead, it has been suggested that the dose metric to quantify nanomaterial uptake can be standardized to enable comparison of results obtained between different studies.^11^

The choice of an appropriate dose metric is critical for results to be relevant to their toxicological effects,^12^ yet proper determination is convoluted in the case of nanomaterials and has been subject to debate.^13^ Mass is the most widely reported metric for toxicological studies due to its linear relationship with small molecule concentration, yet it fails to account for significant interactions resulting from nanomaterial morphology, surface area, chemistry, *etc*.^12^ Moreover, mass comparisons between nanomaterials with dissimilar properties such as size or density are challenging to interpret. Alternative dose metrics have been developed to more adequately describe particle quantities with respect to their observed toxicological response.^6^ Particle number concentration (PNC) is a fundamental measurement used to quantify the number of nanomaterial particles per unit volume.^14^ PNC is a standardized metric for delivered dose when quantified in terms of whole cells, subcellular compartments, or cell volume, enabling comparison of results from different studies.^15^ In contrast to mass concentration, PNC accounts for the structural and interactive components of nanomaterials using discrete, fundamental units,^16^ which can additionally be converted into estimated physical quantities such as surface area concentration. Therefore, PNC can be a useful metric to standardize the quantification of nanomaterial uptake.

Despite its analytical value, PNC measurements in biological studies can be technically challenging to obtain for certain nanomaterials as suitable experimental methods depend on individual measurable properties.^8,17^ The most common approach is direct imaging and counting of internalized particles using high resolution microscopy techniques. Optical microscopies such as confocal fluorescence are easily accessible and can produce 3-dimensional representations which can be useful for per-cell PNC, but particle sizes must be larger than resolution limitations.^18^ Super-resolution microscopy techniques improve the lateral resolution but still remain limited for smaller particle sizes and often require labeling with specialized fluorophores.^8^ Transmission electron microscopy (TEM) offers superior lateral resolution and additionally can distinguish organelles, such as endosomes, to enable per-organelle PNC.^19^ However, the contrast of most non-electron-dense nanomaterials in TEM is limited when imaging in cells stained with heavy atoms. Additionally, quantification often requires a substantial number of sections to image a large enough volume,^8^ or complex techniques such as ion beam milling^20^. Other techniques have been developed to improve upon various limitations by modeling experimental data using statistical analyses.^16, 21, 22^ This combined approach has demonstrated a substantial ability to investigate complex topics, including the random probability distribution of quantitative uptake^21^ and subsequent heterogeneity between endosomes.^22^ Thus, the use of mathematical modeling can further improve the accuracy of these quantitative experimental methodologies.^23^

Conventional analytical techniques are often inadequate for characterizing many nanomaterials with unusual properties, such as single-walled carbon nanotubes (SWCNTs),^17, 24^ which are under investigation for various uses in biomedical applications.^25–28^ SWCNTs exhibit intrinsic near-infrared (NIR) fluorescence emission that is photostable and environmentally-sensitive,^29–31^ and are produced as a mixture of species, or chiralities, which can be identified by their chiral indices (*n,m*).^31^ These advantageous properties have been leveraged to achieve multiplexed optical imaging^28, 32–34^ and sensing^25–27, 35, 36^ in addition to drug- and gene-delivery in live cells,^37, 38^ plants,^39, 40^ and animals.^34, 41^ Such uses necessitate their accurate *in situ* characterization in biological systems, however, the 1-dimensional structure generally makes the direct visualization of individual SWCNTs with appropriate resolution difficult. Although single-SWCNT measurement techniques have been developed,^42–46^ their use has generally been limited solutions, on devices, or otherwise adsorbed to a substrate.

An immediate consequence of the lack of nanometrology techniques that function in live cells is that several fundamental gaps exist in our knowledge of nano-bio interactions, for instance, is there an intrinsic limit in the number of SWCNTs that can enter a cell or a single organelle in a given amount of time? Moreover, a considerable number of factors have been shown to impact the biocompatibility and biodistribution of SWCNTs,^47–49^ suggesting that nanotoxicity and relevant characterizations should be assessed independently for each SWCNT formulation.

In this work, we present a ‘hyperspectral counting’ technique to report the absolute number of emissive SWCNTs within live cells and single endosomes. Using diffraction-limited hyperspectral microscopy,^16^ we acquired spatially-defined spectroscopic data of multiple carbon nanotube emission bands, from different SWCNT chiralities, within live cells. We then performed Monte-Carlo simulations to estimate SWCNT counts from the number of ROIs and number of emission peaks detected. Applying this methodology, we discovered that SWCNT uptake is rate-limited by the cell itself. During 30-minutes of incubation, endocytic uptake is limited to approximately 3,000 SWCNTs per cell. Multiple SWCNTs, loaded within single endosomes even at relatively low incubation concentrations, did not result in SWCNT self-interaction or aggregation. The method also identified significant heterogeneity in nanomaterial distribution among endosomes within a given cell. Consequently, single statistical descriptors such as the mean or median number of nanoparticles per endosome are not sufficiently accurate for describing nanotube uptake by cells, which should be considered in terms of distributions instead.^8^ This work presents a method to quantify cellular and local/subcellular concentrations of a heterogeneous nanomaterial, with implications for nanotoxicology, drug/gene delivery, and nanosensor fields.

## Results And Discussion

### Hyperspectral characterization of carbon nanotube aggregates

Our first goal was to investigate the potential for near-infrared hyperspectral microscopy to identify aqueously-dispersed photoluminescent SWCNTs and aggregates thereof. We selected HiPco SWCNTs, non-covalently dispersed via sodium deoxycholate (SDC) as a model nanomaterial. SDC disperses SWCNTs with high efficiency, sufficiently encapsulates the SWCNT sidewall to prevent optical modulation by the chemical environment and does not alter the intrinsic chirality distribution following dispersion.^50^ HiPco SWCNTs were dispersed in SDC via probe-tip ultrasonication for 30 minutes. For experiments involving live cells, free SDC was removed via 100kDa Amicon filtration. The resulting SDC-SWCNT complexes were stable, with a free SDC concentration of ~ 2.4 mM within the critical micelle concentration (CMC) range (2-6 mM).^51^ We found that SDC-SWCNT complexes remained colloidally stable when diluted in 10% serum, despite decreasing the SDC concentration below the CMC (free SDC <0.02 mM). The SWCNTs were internalized by cells via energy-dependent endocytosis, as confirmed by incubating HeLa cells with SDC-SWCNTs at 4°C and 37°C (Fig. S2). Within live cells, stable SWCNT emission was detectable at 6 and 24 hours after initial uptake (Fig. S3). The movement of SWCNTs in the cells was consistent with localization within lysosomes (Movies S1).^52^

To obtain samples that were dispersed under identical conditions but differed in their degree of purification, we varied the centrifugation step (Fig. S1). One sample was centrifuged at 15,000 x *g* for 5 minutes (referred to as the ‘5-minute sample’) and the other was ultra-centrifuged at 250,000 x *g* for 30 minutes (referred to as the ‘30-minute sample’). At both these accelerations, large aggregates of SWCNTs and some carbonaceous impurities sedimented into a pellet, leaving primarily singly-dispersed SWCNTs or aqueously-dispersed nanotube bundles.^50^

We conducted bulk optical characterization of the material to assess the degree of aggregation. Optical absorbance spectroscopy, Raman spectroscopy, and near-infrared (NIR) photoluminescence are three widely used methods, for which documentary standards have been published.^50^ Optical absorbance spectra of the two samples differed, with higher background absorption and significantly lower peak-to-valley ratio for the 5-minute sample (Fig. S4a). This metric reflects the higher carbonaceous impurities in the 5-minute sample but can also result from aggregation. However, no noticeable wavelength shifts or broadening were detected in the E_11_ absorption peaks, which would potentially denote SWCNT-SWCNT contact/bundling (Fig. S4a). The radial breathing mode of the resonant Raman spectrum was identical for both samples (Fig. S4b) and the characteristic aggregation peak (~267 cm^−1^)^53^ was not detected for either suspension. Photoluminescence emission under 730 nm excitation was significantly higher for the 30-minute sample, consistent with the better dispersion observed in the absorption spectrum (Fig. 4c). Lastly, we characterized the chirality-dependent properties with a two-dimensional photoluminescence excitation-emission (PLE) map (Fig. S5). For the 12 chiralities observed, emission peaks were red-shifted slightly, by < 0.5 nm in the 5-minute centrifugation sample relative to the 30-minute centrifugation sample, but excitation peaks, emission width (full-width at half maximum, FWHM) and excitation width (FWHM) did not show statistically-significant differences (Table S1).

Near-infrared photoluminescence microscopy was used to interrogate both samples at the single-ROI level. Dilute concentrations of the two preparations were adsorbed to a glass surface, rinsed, and imaged in aqueous solution at high magnification (100X) under 730 nm excitation using a NIR hyperspectral microscope. Broadband NIR photoluminescence images integrated across the emission range of 900-1500 nm, and hyperspectral cubes^33^ (spatially-resolved emission spectra of each imaged pixel) from 900-1400 nm were obtained from the same field of view (Fig. 1a). The full spectra were acquired for each spatial region-of-interest (ROI) in the entire field-of-view, from which we counted the number of distinct peaks in the emission spectra (Fig. 1b). Each ROI in the hyperspectral cube was represented by colors mapped to the number of emission peaks detected (Fig. 1c).

**Figure 1.**
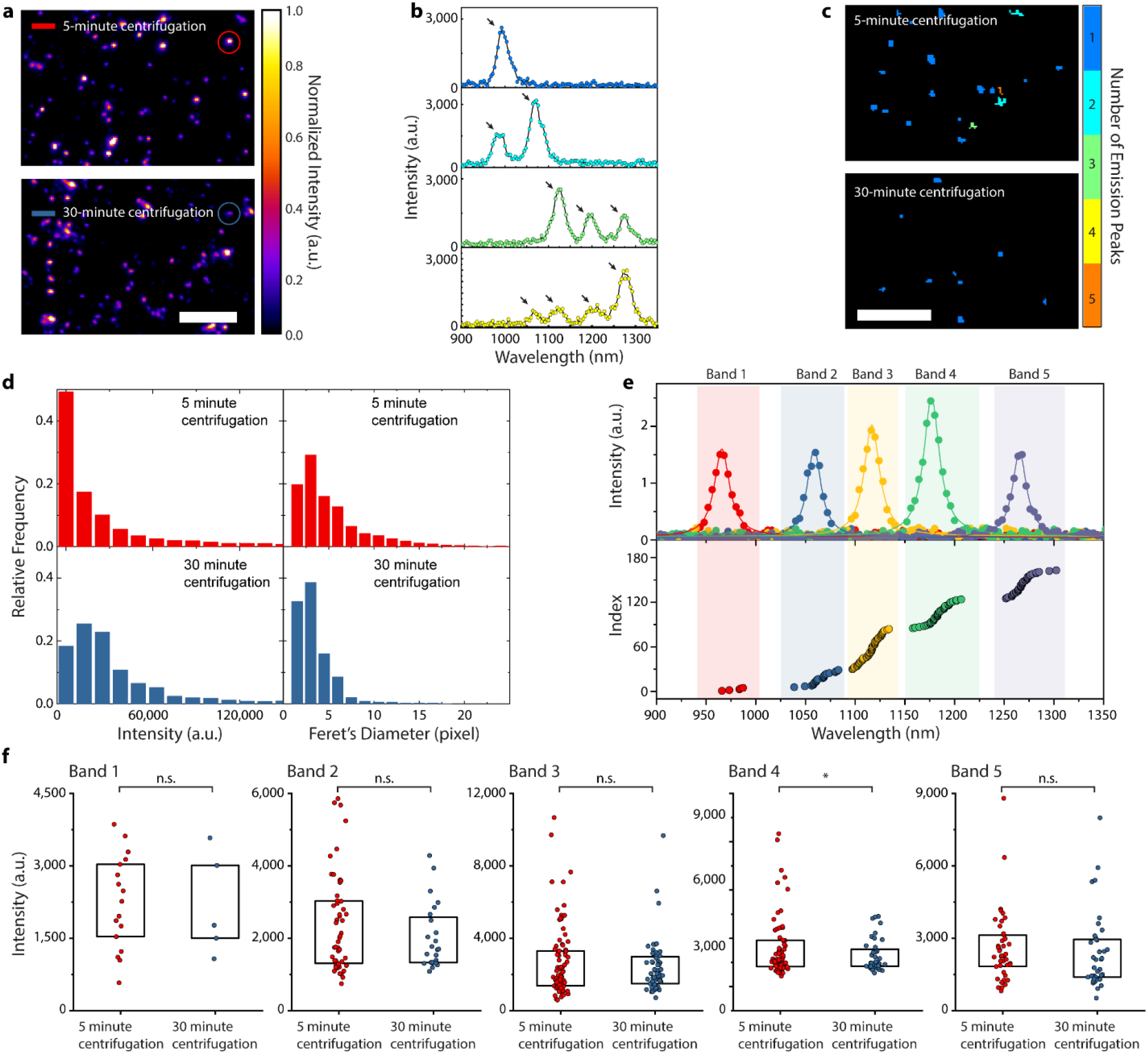
Single-nanotube hyperspectral microscopy of surface-adsorbed SDC-SWCNTs. (a) Broadband near-infrared image of SDC-SWCNTs at 100X magnification. Scale bar = 10 μm. (b) Representative spectra from individual ROIs selected from the 5-minute centrifugation sample, arrows highlight emission peaks. (c) Hyperspectral image of SDC-SWCNTs from each sample at 100X magnification, with each region-of-interest (ROI) false-colored by the number of emission peaks detected. Scale bar = 10 μm. (d) Histogram of total intensity and Feret’s diameter from two SDC-SWCNT sample preparations. (e) Representative spectra (data points fit by Lorentzian functions) from each emission band. Center wavelengths from all SDC-SWCNTs in the 5-minute sample plotted in ascending order. (f) Scatter plot of peak emission intensity of all individual SDC-SWCNTs from the 2 preparations. Boxes represent 25-75% of the data. Statistical comparisons are unpaired t-tests with Welch’s correction.

We assessed quantification of the nanotubes by several methods using the different types of acquired data. We quantified ROIs by brightness and apparent size, as well as by wavelength-defined emission bands. We measured the brightness and apparent size of photoluminescent ROIs, observed from over 2,500 ROIs in each condition (Fig. 1d). The median integrated emission intensity from the 30-minute (centrifugation) sample was approximately 2X higher than the 5-minute sample (19,863 ± 1,243 a.u. vs. 9,073 ± 2,052 a.u.). In contrast, the median Feret’s diameter (which models the size by fitting each ROI to an ellipse) for the 30-minute sample was ~ 30% smaller (2.83 ± 0.045 pixels vs 4.12 ± 0.064). We pooled all individual emission peaks and independently fit each with a Lorentzian function to obtain the peak intensity, center wavelength and FWHM. For the 30-minute sample, the center wavelengths clustered into 5 distinct bands (Fig. 1e) corresponding to chiralities [(8,3), (6,5)], [(7,5), (10,2)], [(9,4), (7,6)], [(12,1), (8,6), (11,3)] and [(10,5), (8,7), (9,5)], respectively (Band edges and bandwidths resulting from the k-means clustering of emission bands are found in Table S2). Center wavelengths for the 5-minute sample also clustered into the same bands (Fig. S6).

Finally, we compared single ROIs after clustering to assess the optical properties within individual bands for the two SWCNT preparations. We measured the peak emission intensities from emission bands within individual ROIs (Fig. 1f). The intensity distributions of both samples were broad, in part because of intrinsic heterogeneity in SWCNT brightness due to factors including length, endohedral content, defect density, surfactant microenvironment, and oxidation state.^50^ In our experimental setup, these factors were further convolved by the unequal excitation efficiency (on-resonance, off-resonance, and k-band phonon absorption for different chiralities) due to single-wavelength excitation.^54^ Although more ROIs with significantly higher intensities were present in the 5-minute sample, no statistically-significant differences between the two samples were observed (except in band 4, p < 0.05). In contrast, emission wavelengths in the 5-minute sample were red-shifted in bands 3, 4 and 5, consistent with the PLE results (Fig. S7, values in Table S3).

### Model to estimate number of carbon nanotubes from emission bands

Because >12 nanotube chiralities were binned into 5 emission bands due to spectral overlap, we asked whether *a priori* knowledge of the emission band distribution of the nanotube sample could be used to accurately approximate the discrete probability distribution of nanotubes per ROI from the probability distribution of emissive peaks. Though we do not know the number of photoluminescent SWCNTs in an ROI, the emission from each SWCNT in that ROI must lie within one of the 5 mutually exclusive bands in wavelength space (Fig. 2a). In a single band, the experimental observation is binary – zero peaks if no emitter is present, and one peak if one or more emitters are present. The full emission spectrum (900 – 1400 nm) of any ROI can therefore at most show 5 distinct peaks.

**Figure 2.**
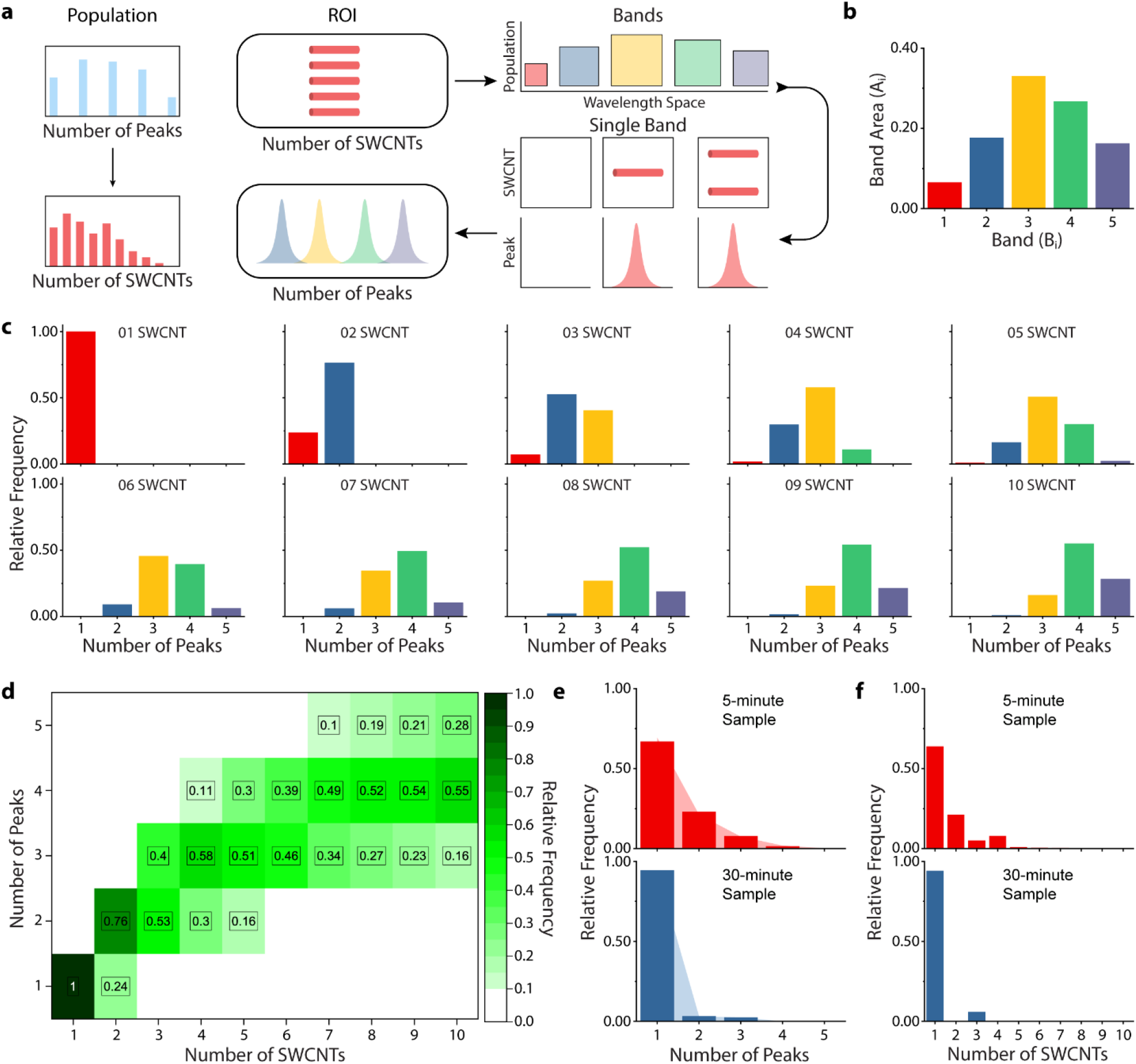
Monte-Carlo model to estimate emissive SWCNTs from emission peaks. (a) Schematic of model to compute the distribution of the number of SWCNTs per ROI from the distribution of the experimentally-measured number of emission peaks per ROI. (b) The population distribution histogram for the 5-minute SWCNT preparation. (c) Relative-frequency histograms representing the number of emission peaks detected for a specific number of SWCNTs for an individual ROI. (d) Heat map of the number of SWCNTs per ROI and the probability distribution of the number of detected emission peaks. Values below 0.1 are not shown, for clarity. (e) Histograms quantifying the experimentally determined relative-frequency of emission peaks from each sample, with shadowed lines representing a quality-of-estimates check from the model output. (f) Relative-frequency histogram of the number of SWCNTs per ROI in the two preparations, as calculated by the model.

We developed a three-step computational method to approximate the number of emissive SWCNTs present in an ROI from the number of emission peaks. First, we formulated a mathematical framework to distribute ‘n’ nanotubes into ‘m’ bands of different sizes. In this extension of the classical combinatorial probability problem of ‘distributing *n* balls in *m* boxes’, the relative size of each box is the relative nanotube population present in each band and can be directly calculated from the experimentally determined chirality distribution of the population. This is an intrinsic property of a carbon nanotube preparation, convolved with the experimental detection parameters of the setup. We determined the band size for the 5-minute sample from the total number of SWCNTs detected via hyperspectral microscopy (Fig. 2b). The distribution was consistent between the 5-minute and 30-minute preparation and matched the results from PLE measurements in solution (Fig. S8).

The probability *ρ*_*i*_ of any carbon nanotube belonging to a specific band *B*_*i*_ in an ROI is:

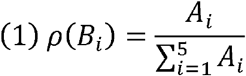

Where *A*_*i*_ correlates to the SWCNT population in *B*_*i*_.

Next, we generated a mapping function for the number of SWCNTs in an ROI and the number of emission peaks detected by solving a two-step process: (1) If an ROI contained *N* SWCNTs in total, where *N* ranges from 1 to 10, how many SWCNTs on average would belong to each band? (2) As only zero or one peak can be observed per band, how many peaks would be detected in the full emission spectrum from that ROI?

We chose 10 as the upper limit for the total number of nanotubes present within an ROI for 2 primary reasons: First, the intensity distribution from individual ROIs (Fig. 1f) was consistent with one broad population likely arising from one emitter, with outliers that were approximately twice as bright. Second, at the concentration range used through this work, the experimentally detected number of emission peaks per wavelength band was consistently less than two. For a system with 5 bands, this limits the number of SWCNT per ROI to ten.

In any band *B*_*i*_, *Φ* is the number of emission peaks detected when *P* SWCNTs are present:

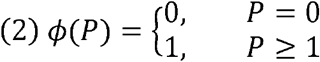

As there is no closed-form analytical solution for this system, we sought a numerical approximation using Monte Carlo simulations. For each N (number of SWCNTs present, ranging from 1 to 10), we ran 5,000 simulations to obtain a histogram of the number of emission peaks detected per ROI (Fig. 2c, details in Supplementary Text 1). These results mapped the number of SWCNTs present in a single ROI to the probability of detecting a specific number of emission peaks (Fig. 2d). For example, if 4 SWCNTs were present in an ROI, the probability of detecting 2 peaks was 0.30, of 3 peaks was 0.58 and 4 peaks was 0.11.

Finally, we extended the single-ROI model to an entire population. Essentially, any system of SWCNT-containing ROIs can be characterized by either a distribution of the number of nanotubes or by a distribution of the number of emission peaks (Fig. 2a). For a population of *m* ROI (*R*_*i*_), where each ROI contained *N*_*m*_ SWCNTs that were detected as *P*_*m*_ peaks, these two sets are equivalent:

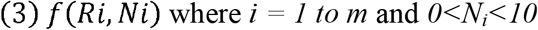

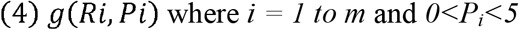

A probability mass function from one variable mapped into a probability mass function from the other i.e., for a population containing ROIs with *N*_*j*_ SWCNTs and *P*_*i*_ observable peaks:

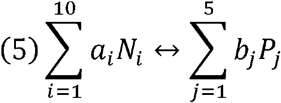

The relative probabilities associated with the number of SWCNTs (*a*_*i*_) and the number of peaks (*b*_*j*_) summed to 1:

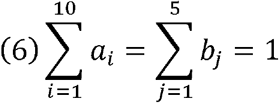

The number of emission peaks per ROI (coefficients *b*_*j*_) were directly calculated from hyperspectral data of the two surface-adsorbed SDC-SWCNT samples (Fig. 2e). Only ~ 70% of ROIs in the 5-minute sample had one emission peak, in contrast to ~95% for the 30-minute sample. Using the *b*_*j*_ coefficients as inputs, we solved the system of linear equations in (5) and (6) for the unknown coefficients *a*_*i*_ via the least-squares method to obtain a distribution of the number of SWCNTs per ROI (Fig. 2f). To test the quality of the solution, the coefficients *a*_*i*_ were used to generate a *b*_*j*_’ and directly compared with the experimentally determined *b*_*j*_. The parameters obtained regenerate a distribution for the number of emission peaks with reasonable agreement with the experimental data (shaded lines in Fig. 2e, with experimental data represented by solid bars).

### Endocytic uptake results in multiple carbon nanotubes per endosome

We introduced SWCNTs to live cells under short incubation times to investigate endosomal accumulation. Hyperspectral microscopy was used to quantify SDC-SWCNT uptake in live mammalian cells, with the primary goal of extracting quantitative parameters that could be objectively compared across multiple experiments. Our model system was defined as SWCNT uptake via a 30-minute pulse in HeLa cells, a timepoint which results in nearly complete uptake of cell-associated nanotubes but before reverse trafficking of these SWCNT-containing endosomes is initiated.^55^ The 30-minute preparation was used to ensure that the SDC-SWCNT sample itself was dispersed well with minimal aggregation. HeLa cells were incubated for 30 minutes with the 30-minute SDC-SWCNT preparation in cell media, at concentrations spanning two orders of magnitude from 0.1 – 10 mg/L. Cells were thoroughly washed to remove unbound SWCNTs and placed at 4°C for 30 minutes in fresh media to reduce movement before imaging. For each cell, a z-stack of NIR broadband fluorescence images through the entire volume and a hyperspectral cube at the central z-position were sequentially acquired. Within this acquisition time (< 2 minutes), there was minimal movement of either the cell or the ROIs. The photoluminescence images of SWCNT emission from HeLa cells were consistent with SWCNTs bound to the cell membrane, on either the outside of or just internalized into the cell (Fig. S9). In our experimental setup, we previously showed that the presence of relatively dim SWCNTs that were not detected is negligible,^56^ *i.e.* almost all ROIs with NIR emission were present in the photoluminescence image.

We quantified SWCNTs within endosomes by several methods. The total number of ROIs within each cell were counted from the maximum intensity projection image. At 30-minute incubation, these ROIs were primarily early endosomes.^55^ Most ROIs contained just one emission peak at 0.1 mg/L, but the number of peaks ranged from one to five after incubating with 1, 5 and 10 mg/L of SWCNTs (Fig. 3a). These results are direct evidence of multiple SWCNTs within each ROI. Surprisingly, the photoluminescence intensity did not reflect this heterogeneity, as the emission intensity from individual ROIs at 1, 5 and 10 mg/L was not statistically different, for any of the emission bands (Fig. 3b). This finding indicates that emission intensity itself was an unreliable metric for quantifying carbon nanotube uptake. The emission wavelengths also did not exhibit any consistent modulation as a function of SDC-SWCNT concentration (Fig. S10), indicating no notable SWCNT-SWCNT interactions.

**Figure 3.**
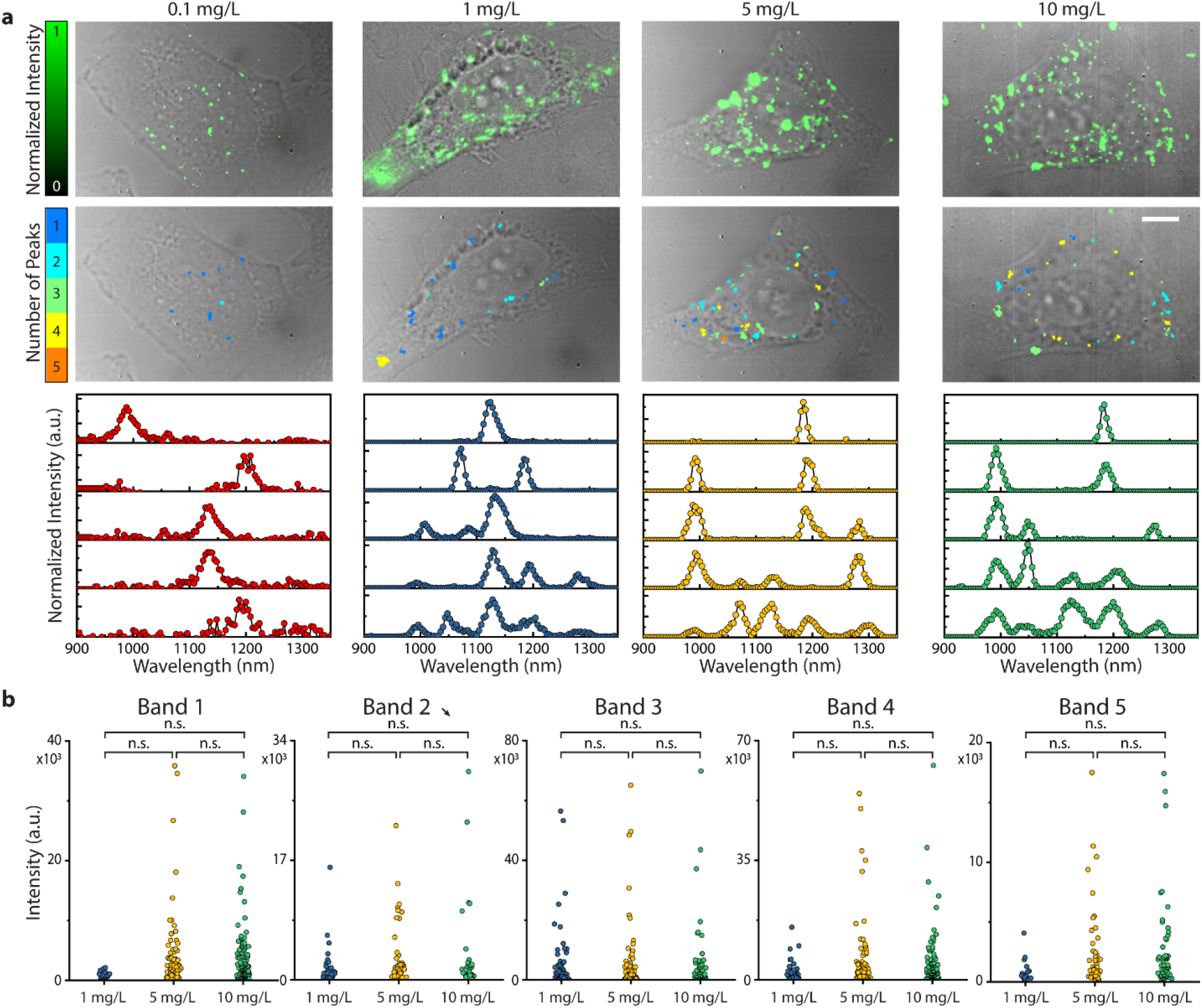
Near-infrared hyperspectral microscopy of SWCNT uptake in HeLa cells. (a) Broadband fluorescence maximum intensity projection image, computed image with each ROI false-colored by the number of emission peaks detected and representative spectra from individual ROIs, at 0.1, 1, 5 and 10 mg/L SDC-SWCNT loading concentration. Scale bar = 10 μm. (b) Intensity of individual ROIs for each loading concentration. One-way ANOVA was performed using Holm-Sidak’s multiple comparison test.

### Saturation of nanotube uptake in cells and endosomes

Particle number concentration measurements using the combined hyperspectral and computation counting technique were performed to quantify the concentration-dependent cellular uptake and sub-cellular distribution of single-walled carbon nanotubes. The absolute count of the SWCNT-containing endosomes within a cell was experimentally determined via high-magnification live-cell fluorescence microscopy (as shown in Fig. 3a). Normalized by the projected area of each cell, we assessed the number of SWCNT-containing ROIs per unit area as a function of SWCNT-loading concentration (Fig. 4a). The areal density of SWCNT-containing ROIs increased with the SWCNT concentration administered to the cells (Spearman correlation = 0.90 with p < 0.0001) and was accurately described by an extended Langmuir adsorption model (R^2^ = 0.996), plateauing at ~ 0.29 ROI per μm^2^. The data at 0.1, 1 and 5 mg/L were statistically different from each other, but no significant differences were observed between the values at 5 and 10 mg/L (gray shaded box in Fig. 4a).

**Figure 4.**
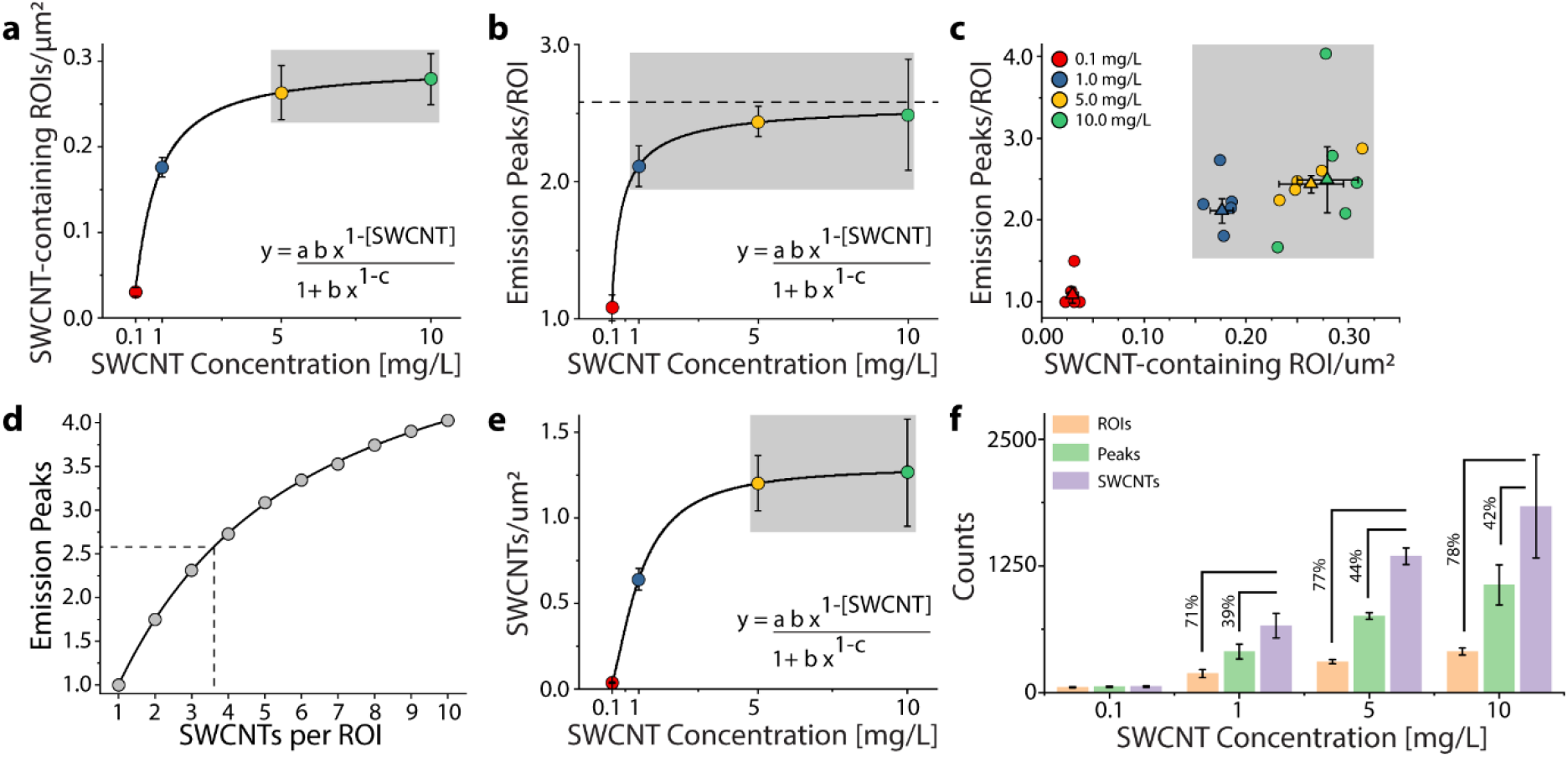
Multiparameter characterization of SWCNT uptake in HeLa cells. (a) Density of SWCNT-containing ROIs as a function of loading concentration. Line is a fit of the Langmuir isotherm equation to the data; error bars denote SEM. Gray region did not show a statistically significant difference. (b) Average number of emission peaks per ROI, as a function of loading concentration. Line is a fit of the Langmuir isotherm equation to the data; error bars denote SEM. Gray region did not show a statistically significant difference. (c) Scatter plot of the emission peaks per ROI vs. density of SWCNT-containing ROIs. Individual cells are circles; triangles represent the mean. Errors bars denote SEM. Gray regions could not be separated via k-means clustering. (d) Mapping between the number of emission peaks detected within one ROI and the computed number of emissive SWCNTs. Dashed lines correspond to the limiting value of the number of emission peaks per ROI, determined by the fit (dashed line) in panel b. (e) Density of SWCNTs as a function of loading concentration. Line is a fit of the Langmuir isotherm equation to the data. Error bars denote SEM. Gray region did not show a statistically significant difference. (f) Comparison of absolute SWCNT counts as a function of loading concentration, using 3 metrics. Error bars denote SEM. Percentage differences calculated from the SWCNT count.

The number of SWCNTs per endosome was experimentally determined for each cell using hyperspectral microscopy. For each spatially distinct ROI, we directly counted the number of distinct emission peaks in the 900-1400 nm wavelength rang. The mean number of emission peaks per ROI increased with SDC-SWCNT concentration (Spearman correlation = 0.68 with p < 0.001) and was accurately described by an extended Langmuir adsorption isotherm (R^2^ = 0.99,), plateauing at ~ 2.58 emission peaks/ROI (Fig. 4b). Except for the data at 0.1 mg/L SWCNT loading concentration, no statistically significant differences were observed between the data at 1, 5 and 10 mg/L (gray box in Fig. 4b). A scatter plot of the density of SWCNT-containing endosomes per cell and the number of distinct emission peaks per endosome (Fig. 4c) revealed a high degree of correlation (r = 0.80, p<0.0001, paired t-test). Although the data at 0.1 and 1.0 mg/L appeared distinguishable from 5 and 10 mg/L, an unbiased k-means clustering analysis was only able to accurately separate the 0.1 mg/L data from the higher concentrations (Fig. S11). Cells incubated with 1 mg/L could not be identified from cells incubated with 5 and 10 mg/L SWCNT (gray region in Fig. 4c could not be separated). Combined, these results indicate that the density of SWCNT-containing ROI saturate above 5 mg/L loading concentration, while the number of emissive peaks per ROI plateaus by 1 mg/L loading concentration.

Using the computational model, we calculated the emissive SWCNTs within each ROI from the number of distinct emission peaks. The nanotube band distribution of the 30-minute SDC-SWCNT sample in cells, obtained using the same hyperspectral analyses used for Fig. 2b, was significantly different from the solution measurement (Fig. S12). This result likely arose from the known chirality-dependent modulations in SWCNT emission wavelength and intensity by the intracellular environment.^49, 57^ Following the procedure developed in a previous section for SDC-SWCNT adsorbed on a surface, we obtained an analogous heat map of the number of SWCNTs per ROI and the probability distribution of the number of detected emission peaks for SDC-SWCNTs in cells (Fig. S13). Our analysis generated a calibration curve between the experimentally detected number of emission peaks in an ROI and the least-squares estimate of the number of emissive SWCNTs physically present (Fig. 4d). The density of emissive SWCNTs per cell (Fig. 4e) increased with concentration (Spearman correlation = 0.81 with p < 0.0001) and plateaued at ~ 1.3 SWCNTs per μm^2^ (Langmuir fit, R^2^ = 0.99). No statistically significant differences were detected between the two highest concentrations. For a 30-minute incubation of SDC-SWCNTs in HeLa cells, we found the linear uptake regime to be below the 1 mg/L SDC-SWCNT concentration range in media.

We compared the number of SWCNT-containing ROIs, the total number of emission peaks and the particle number concentration to quantify nanotube uptake within a cell (Fig. 4f). Assuming the SWCNT count as the reference standard, counting the total number of emission peaks systematically underestimated the actual values by ~ 40%, while counting the ROIs underestimated the actual counts by ~ 70%. The mean nanotube signal per cell, as quantified by photoluminescence intensity in broadband images, was the least accurate, undercounting the SWCNT concentration by 15-fold (Fig. S14). At the highest loading concentration of 10 mg/L, an average cell contained 406 ± 35 ROIs, 1062 ± 198 emission peaks, and 1838 ± 509 emissive nanotubes. As approximately 1/3^rd^ SWCNTs in the HiPco sample are non-emissive (metallic and semi-metallic), we scaled the number of emissive SDC-SWCNTs by 1.5 to calculate the total number of SWCNTs present.

Using the particle numbers above, we obtained a quantitative description of SWCNT partitioning into individual cells and endosomes (detailed calculations in Supplementary Text 2). For an SDC-SWCNT loading concentration of 10 mg/L (~ 130 nM), approximately 3,000 SWCNTs were endocytosed per cell, with an average of 4 SWCNTs per endosome. The average HeLa cell is 3,000 μm^3^ in volume and the typical endosome is ~250 nm in diameter.^58^ This corresponds to a SWCNT concentration of ~2 nM within a cell, indicating an effective partitioning of 1.5% of the SWCNT concentration in solution into a cell. However, the SWCNT concentration within the endosomes is ~300 nM, which is 2.3 times the concentration in solution.

### Quantifying intercellular and intracellular heterogeneity

We assessed inter- and intra-cellular heterogeneity of SWCNT uptake and distribution. We obtained population statistics by pooling data from individual ROIs across multiple cells. From histograms of SWCNT emission peaks per ROI, we found that over 70% of the ROIs at 1, 5 and 10 mg/L contained more than one nanotube (Fig. 5a). The distribution shifted to a higher number of peaks with increasing loading concentration. Analysis of individual cells revealed significant inter and intra-cellular heterogeneity, with minimal dependence of either distribution on SWCNT loading concentrations above 1 mg/L(Fig. 5b). Though multiple factors determine SWCNT uptake by a cell, the specific number of particles associated with each cell is random.^22^

**Figure 5.**
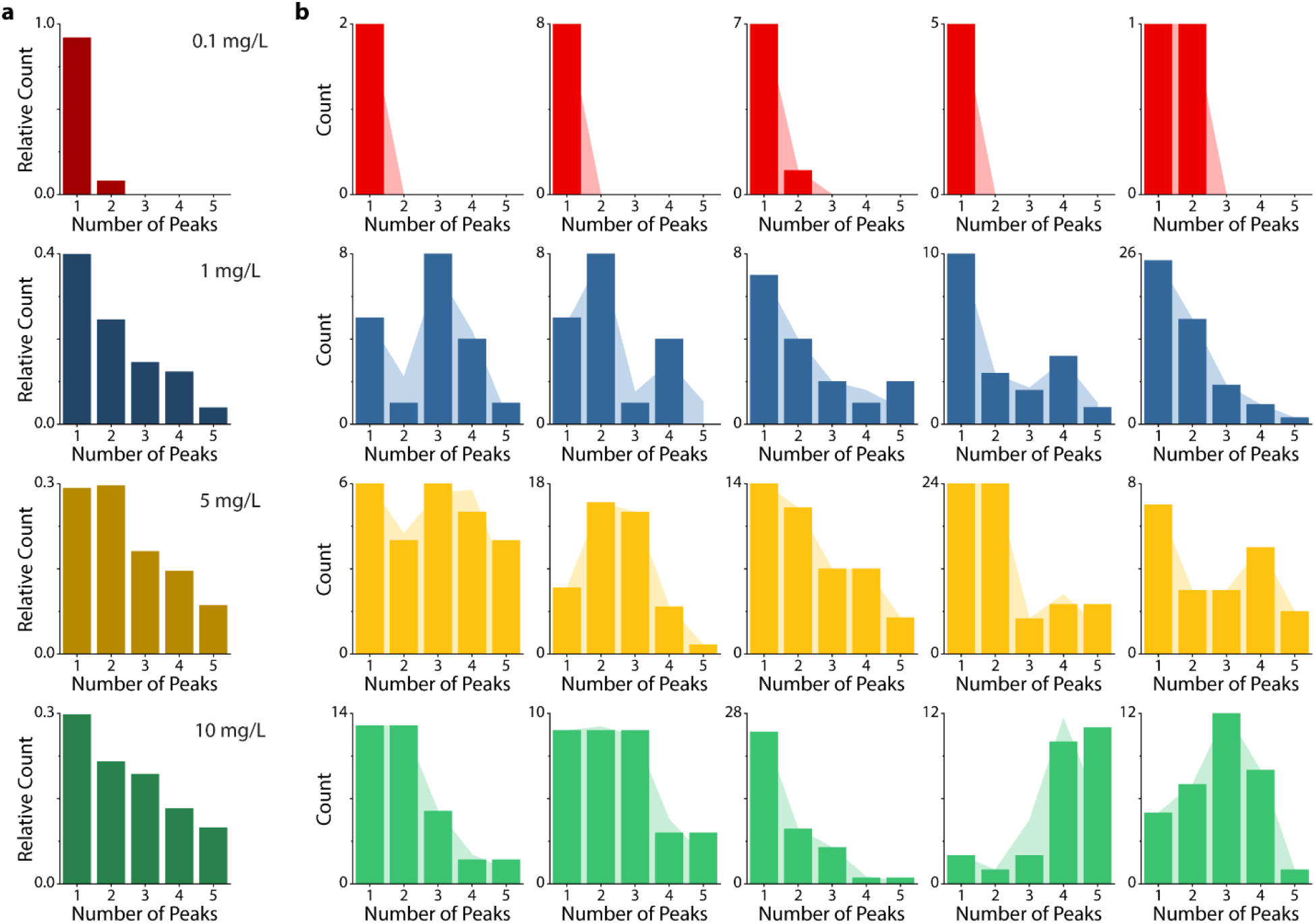
Inter and intra-cellular heterogeneity in SWCNT uptake by HeLa cells. (a) Relative-frequency histogram of the emission peaks detected for the entire cell population, at each SWCNT concentration. (b) Frequency histograms (absolute counts) quantifying the experimentally determined relative-frequency of emission peaks from each cell, with shadowed lines representing a quality-of-estimates check from the model output

In contrast to single homogeneous system comprised of multiple fields-of-view of SDC-SWCNT adsorbed on a surface, the intercellular heterogeneity observed required us to consider each cell to be an independent system. Computationally, this meant solving a separate system of linear equations for each cell, where the experimentally determined distribution of emission peaks per ROI were used to obtain the least-squares estimates of the SWCNT distribution. The corresponding distribution of the number of emission peaks was subsequently calculated to the quality of fit (Fig. 5b, following the same procedure described in Supplementary Text 2, and used to generated Fig. 2e). The number of emission peaks per ROI varied broadly from 1 to 5 within a single cell, at all concentrations above 0.1 mg/L. Additionally, the distribution was also heterogeneous across cells at each concentration, with varying minimum, maximum, and median values. A direct consequence of this heterogenous distribution is that statistical descriptors such as the mean or median number of emission peaks per ROI at any loading concentration only accurately describe a small fraction of the total ROI population.

## Conclusions

In this work, we have developed a nanometrology technique to quantify the uptake of single emitting nanomaterials in living cells. Using NIR hyperspectral imaging, we quantified spectral bands to enable the counting of single SWCNT emitters within single endosomes. We employed experimentally-guided Monte Carlo simulations to further improve the robustness of the method. HeLa cells were determined to internalize ~3,000 SWCNTs when dosed for 30 minutes at a concentration of 10 mg/L, with an average of 4 SWCNTs per single endosome. Our analysis further determined that SWCNT uptake is rate-limited by cells with both the SWCNT-containing endosomes and number of SWCNTs per endosome plateauing at less than 5 mg/L concentration of SWCNTs in media, with a linear uptake regime below 1 mg/L. Moreover, both the intracellular and intercellular distribution of SWCNTs per endosome are highly heterogeneous. The tails of such distributions are significant, as several mechanisms of nanoparticle-induced toxicity result in signaling from individual organelles or cells. Even if the average endosome per cell has between 1-2 nanotubes, the presence of larger quantities in a single endosome could induce localized toxic effects, in addition to generating systematic errors in sensing, imaging and delivery applications. For future applications, calculations of cellular and endosomal nanotube concentrations need to be considered in terms of distributions instead of single statistical descriptors.

With multiple families of fluorophores under development, and hyperspectral microscopy becoming increasingly available in research labs, the framework developed in this work has broad applicability for various nanomaterials. Moreover, the ability to compare uptake of SWCNTs with other types of nanomaterials is possible using particle number concentration (PNC) as the dose metric, promoting advancements in our understanding of complex interactions vital to nanomedicine.

## Methods

### Preparation of Single-Walled Carbon Nanotube Suspensions

Single-walled carbon nanotubes produced by the HiPco process (Unidym) were suspended by probe-tip ultrasonication (Sonics & Materials, Inc.) of 20 mg sodium deoxycholate (SDC) with 1 mg of ‘raw’ SWCNTs in 1 mL of deionized water for 30 minutes at 40% of the maximum amplitude (~ 9 Watts). Following sonication, the dispersions were ultracentrifuged (Sorvall Discovery 90SE) for 30 minutes at 280,000 x *g*. The top ¾ of the resulting supernatant was collected. Concentration was determined with a UV/Vis/NIR spectrophotometer (Jasco) using the extinction coefficient A_910_ = 0.02554 L·mg^−1^ cm^−1^.^33^ To remove free SDC, 100 kDa Amicon centrifuge filters (Millipore) were used to concentrate the nanotube dispersions and re-suspend via mixing by pipette. Nanotubes were prepared immediately before addition to cell media. A photoluminescence excitation-emission contour plot was constructed to identify the nanotube chiralities present in the sample using a custom-built instrument.^59, 60^

### Cell Culture

HeLa CCL-2 cells (ATCC) were grown under standard conditions at 37°C and 5% CO_2_ in sterile-filtered DMEM with 10% heat-inactivated FBS, 2.5% HEPES, 1% glutamine, and 1% penicillin/streptomycin (all Gibco). Cells were plated onto T-75 flasks at 20% confluence and passaged every 3 days. For imaging experiments, cells were plated onto glass-bottom petri dishes (MatTek) and used at 70-80% confluence.

### Hyperspectral Imaging

As described previously,^33^ an instrument to conduct NIR fluorescence hyperspectral microscopy was used to obtain spectrally-resolved images of emissive nanotubes in HeLa cells (Photon etc.). Briefly, a continuous wave (CW) 730 nm diode laser (with output of 230 mW, measured at the sample) was injected into a multimode fiber to produce the excitation source for photoluminescence experiments. A long pass dichroic mirror with a cut-on wavelength of 880 nm was aligned to reflect the laser into an Olympus IX-71 inverted microscope (with internal optics modified for near infrared transmission) equipped with a 100X (UAPON100XOTIRF, NA=1.49) oil-immersion objective (Olympus). Emission from the nanotubes was spatially and spectrally resolved by passing through a volume Bragg grating and into a thermo-electrically cooled 256 x 320 pixel InGaAs array detector. A continuous stack (hyperspectral cube) of 126 spectrally-defined images was obtained between 900 to 1400 nm, collected in 4 nm steps. The data was processed to produce a near-infrared spectrum for every pixel of the image. To quantify the absolute number of nanotubes per cell, z-stacks were constructed from HeLa cells incubated with varying concentrations of SDC-HiPco for 30 minutes, washed with fresh media, and then placed in 4°C for 15 minutes prior to imaging (to temporarily halt endosomal movement). The number of distinct nanotube-containing endosomes was determined by manually counting the z-stack images.

### Two-dimensional excitation/emission photoluminescence plots

Photoluminescence (PL) plots were acquired using a home-built apparatus consisting of a tunable white light laser source, inverted microscope, and InGaAs NIR detector. A SuperK EXTREME supercontinuum white light laser source (NKT Photonics) was used with a VARIA variable bandpass filter accessory to tune the output from 500 – 825 nm with a bandwidth of 20 nm. A longpass dichroic mirror (900 nm) was used to filter the excitation beam. The light path was shaped and fed into the back of an inverted IX-71 microscope (Olympus) where it passed through a 20x NIR objective (Olympus) and illuminated a 200 μL nanotube sample in a 96-well plate (Greiner). Emission from the nanotube sample was collected through the 20x objective and passed through a dichroic mirror (875 nm, Semrock). The light was f/# matched to the spectrometer using several lenses and injected into an Isoplane spectrograph (Princeton Instruments) with a slit width of 410 μm which dispersed the emission using an 86 g/mm grating with 950 nm blaze wavelength. The light was collected by a PIoNIR InGaAs 640 x 512 pixel array (Princeton Instruments).

Excitation, emission, and wavelength corrections and calibrations were performed as follows. The power at each excitation wavelength was measured at the objective with a PM100D power meter (Thorlabs) from which a power spectrum was constructed and used to correct the emission intensities for non-uniform excitation. A HL-3-CAL-EXT halogen calibration light source (Ocean Optics) was used to correct for wavelength-dependent features in the emission intensity arising from the spectrometer, detector, and other optics. A Hg/Ne pencil style calibration lamp (Newport) was used to calibrate spectrometer wavelength.

Acquisition was conducted in automated fashion controlled by Labview code which iteratively increased the excitation laser source from 500 – 824 nm in steps of 3 nm, acquired data with an exposure time of 0.3 seconds for a nanotube concentration of 1 mg/L SDC-HiPco, and saved the data in ASCII format. The spectral range was 930 – 1369 nm with a resolution of ~0.7 nm.

Background subtraction was conducted using a well filled with DI H_2_O. Following acquisition, the data was processed with MATLAB code which applied the aforementioned spectral corrections, created the contours with a Gaussian smoothing function, and constructed figures to be used for manual assignment of nanotube chiralities from the two-dimensional peaks.

### Solution Raman Spectroscopy

All Raman scans and measurements were performed with a Renishaw InVia Raman microscope (Renishaw, Hoffman Estates, IL) equipped with a 785 nm diode laser (300 mW cm^−2^) and a 1 in. charge-coupled device detector with a spectral resolution of 1.07 cm^−1^. Raman spectra were acquired through a 5× objective (Leica, Buffalo Grove, IL), where laser output at the objective was measured to be 100 mW cm^−2^ using a hand-held laser power meter (Edmund Optics, Inc., Barrington, NJ), as previously described.^61^ Data analysis of the spectral images was performed in MATLAB (R2014b) and PLS Toolbox v.8.0 (Eigenvector Research, Inc., Wenatchee, WA). For displayed SERS intensities, baseline subtraction was performed on the collected spectra using a Whittaker filter with λ = 200 cm^−1^.

## Acknowledgements

This work was supported in part (in the D.A.H. laboratory) by the NIH New Innovator Award (DP2-HD075698), NCI (R01-CA215719), the Cancer Center Support Grant (P30 CA008748), the National Science Foundation CAREER Award (1752506), the Honorable Tina Brozman Foundation for Ovarian Cancer Research, the American Cancer Society Research Scholar Grant (GC230452), the Pershing Square Sohn Cancer Research Alliance, the Expect Miracles Foundation–Financial Services Against Cancer, the Cycle for Survival’s Equinox Innovation Award in Rare Cancers, Mr. William H. Goodwin and Mrs. Alice Goodwin and the Commonwealth Foundation for Cancer Research, the Experimental Therapeutics Center, the Kelli Auletta Fund, and the Center for Molecular Imaging and Nanotechnology of Memorial Sloan Kettering Cancer Center, Functional Genomics Initiative, the Alan and Sandra Gerry Metastasis, and Tumor Ecosystems Center. C.C. was supposed by the Eunice Kennedy Shriver National Institute of Child Health & Human Development of the NIH Award F31HD105405. The D.R. laboratory was supported by National Science Foundation CAREER Award # 1844536 and the University of Rhode Island College of Engineering.

## Competing interests

D.A.H. is co-founder and officer with an equity interest in Goldilocks Therapeutics Inc., Lime Therapeutics Inc., and Nirova Biosense Inc. and a member of the scientific advisory boards of Concarlo Holdings LLC, Nanorobotics Inc., and Mediphage Bioceuticals Inc. P.V.J. is a co-founder with an equity interest in Lime Therapeutics, Inc.

