## Supporting Information for "Hyperspectral Counting of Multiplexed Nanoparticle Emitters in Single Cells and Organelles"

### Table of Contents


| | | |
|---|---|---|
| **Figure S1.** | Scheme for preparing single-walled carbon nanotube (SWCNT) samples with differing degrees of dispersion. | S3 |
| **Figure S2.** | Uptake study of SDC-SWCNT complexes by HeLa cells at 4°C and 37°C. | S4 |
| **Figure S3.** | Photostability of SDC-SWCNT complexes in HeLa cells at 6 hours and 24 hours after uptake. | S5 |
| **Figure S4.** | Ensemble optical measurements of the two SDC-SWCNT preparations. | S6 |
| **Figure S5.** | Two-dimensional photoluminescence excitation emission plots of the 5-minute sample and the 30-minute sample. | S7 |
| **Figure S6.** | Clustering of emission center wavelength of individual surface-adsorbed ROIs from both samples. | S8 |
| **Figure S7.** | Emission center wavelength of surface-adsorbed SDC-SWCNT complexes in each band from both samples. | S9 |
| **Figure S9.** | Photoluminescence imaging study of SDC-SWCNT distribution in HeLa cells. | S11 |

| | | |
|---|---|---|
| **Figure S10.** | Emission wavelengths of SDC-SWCNT complexes in HeLa cells at 3 SDC-SWCNT loading concentrations. | S12 |
| **Figure S11.** | K-means analysis of concentration-dependent peaks per ROI vs. ROIs per $\mu m^2$ for single cells. | S13 |
| **Figure S12.** | Population distribution of SDC-SWCNTs in solution and in HeLa cells. | S14 |
| **Figure S13.** | Heat map of the number of SWCNTs per ROI and the probability distribution of the number of emission peaks detected per ROI. | S15 |
| **Figure S14.** | Comparison of four different metrics to quantify SWCNTs in cells. | S16 |
| **Table S1.** | Optical Parameters from PLE Plots | S17 |
| **Table S2.** | Band edges and widths from k-means clustering of emission bands | S22 |
| **Table S3.** | Optical parameters of SDC-SWCNT bands on surface | S23 |
| **Movies S1** | SDC-SWCNT in cells at 6h and 24h | |
| **Supplementary Text 1** | Details of computational model developed | S24 |
| **Supplementary Text 2** | Detailed calculation of nanotube concentration within single cells and single endosomes | S27 |

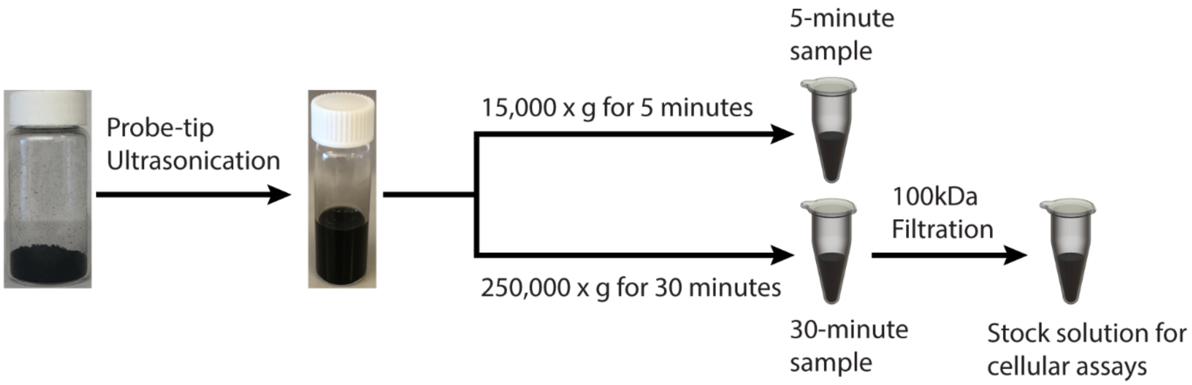


Figure S1. Scheme for preparing single-walled carbon nanotube (SWCNT) samples with differing degrees of dispersion. The “5-minute sample” and the “30-minute sample” were used for cell-free assays. Unbound sodium deoxycholate (SDC) was removed from the 30-minute sample immediately prior to experiments in live cells.

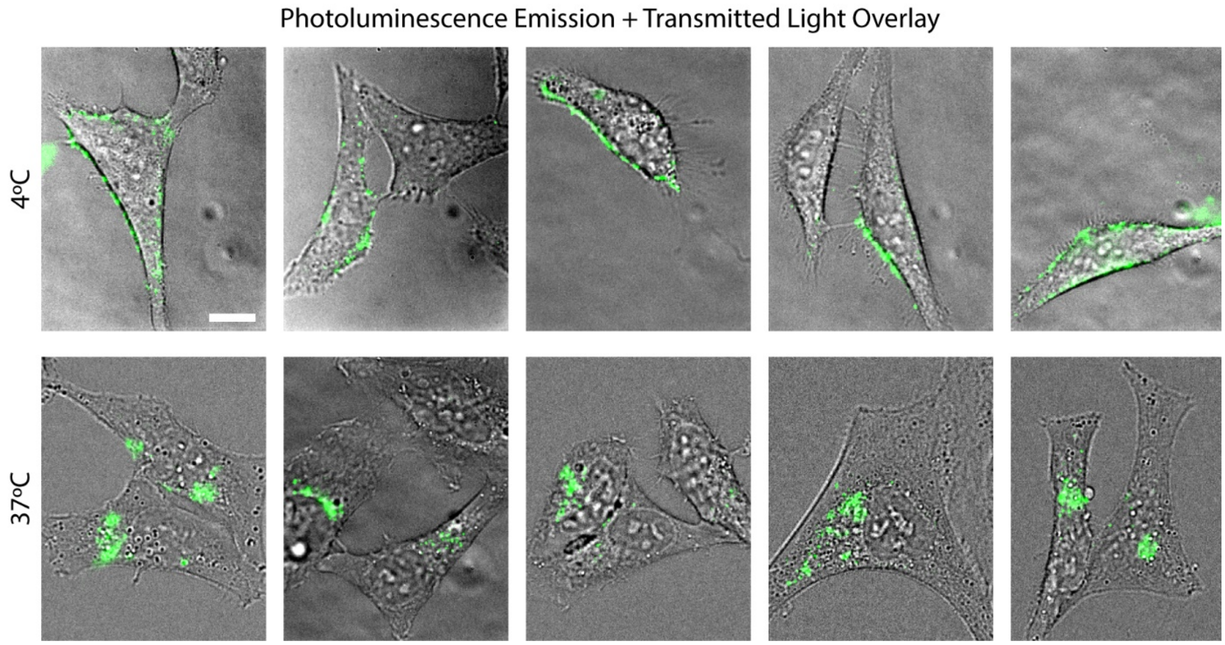


Figure S2. Uptake study of SDC-SWCNT complexes by HeLa cells at 4°C and 37°C. Overlay of transmitted light and near-infrared (NIR) photoluminescence images taken 30 minutes after incubating HeLa cells with 1 mg/L of the SDC-SWCNTs (stock solution for cellular assays) in cell media at 4° C and 37° C. Scale bar is 10 μm.

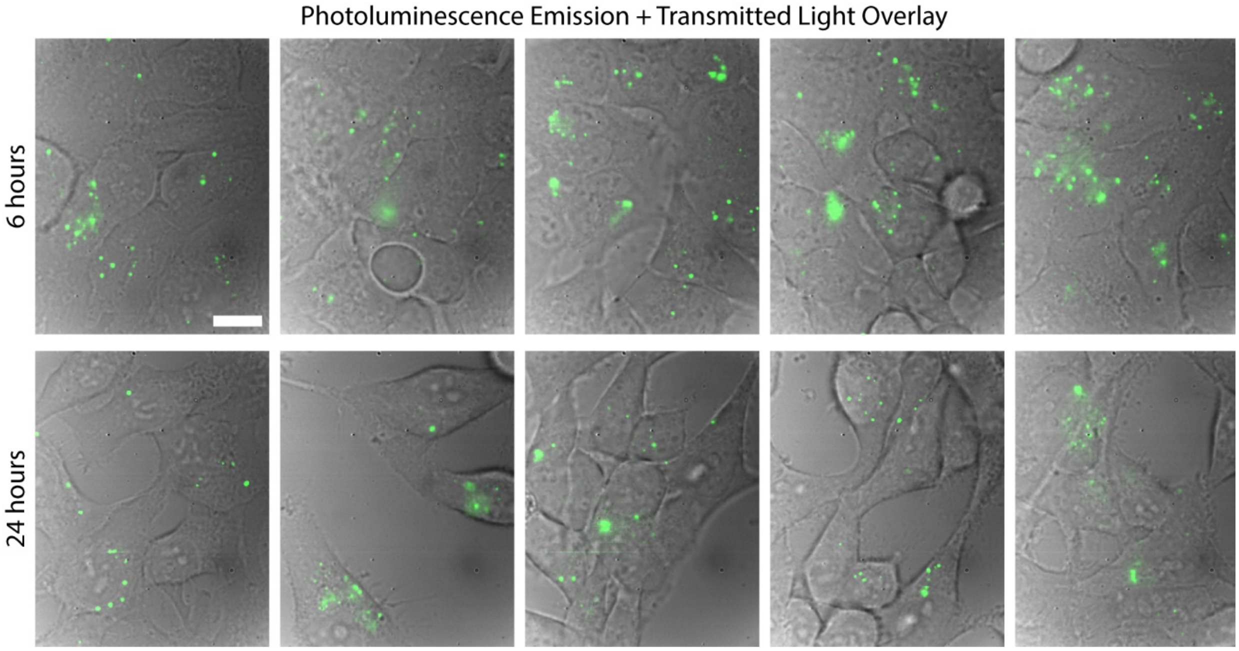


Figure 3. Photostability of SDC-SWCNT complexes in HeLa cells at 6 hours and 24 hours after uptake. Overlay of transmitted light and NIR photoluminescence images of HeLa cells, taken 6 hours and 24 hours after incubation for 30 minutes with 1 mg/L SDC-SWCNTs. Scale bar is 10 μm.

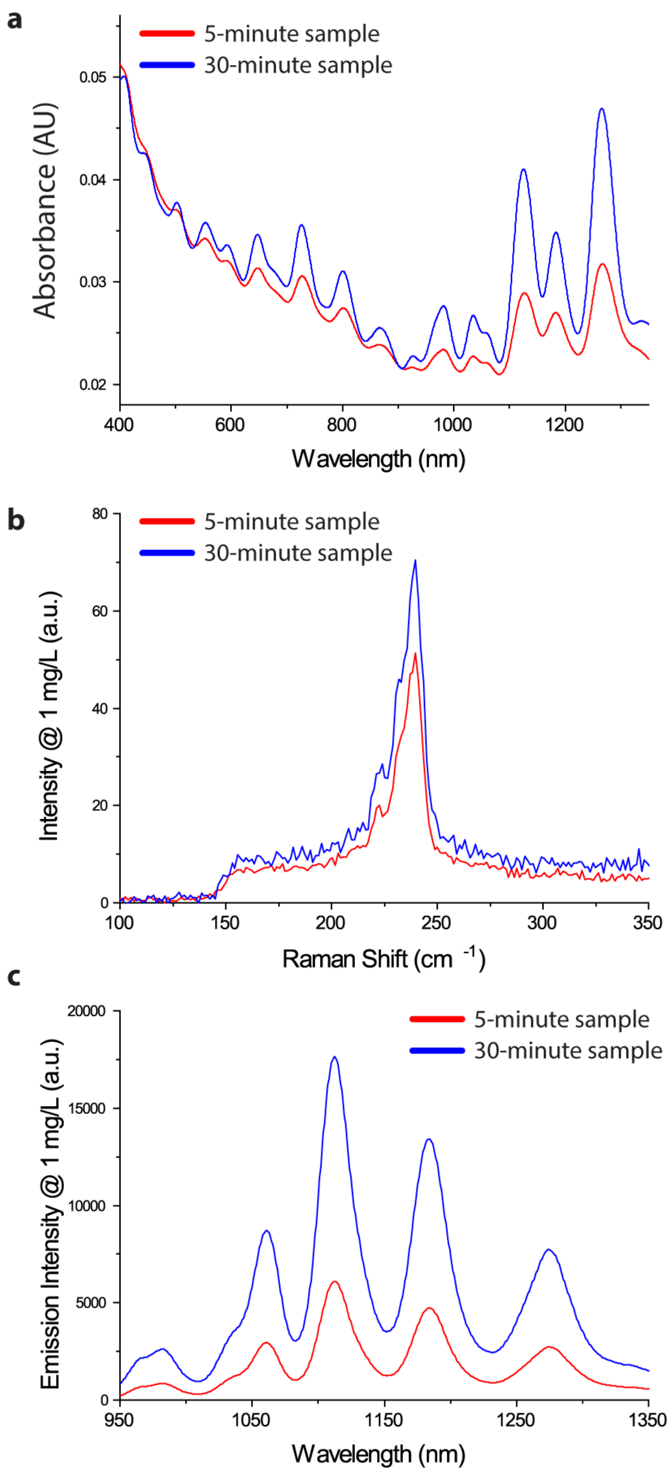


Figure S4. Ensemble optical measurements of the two SDC-SWCNT preparations. (a) Absorption spectra of the 5-minute and 30-minute samples, normalized at 910 nm (corresponding to 1 mg/L effective concentration). (b) Raman spectra of the two samples (1 mg/L) under 785 nm excitation. (c) Photoluminescence emission spectra of the two samples at 1 mg/L concentration under excitation at 730 nm.

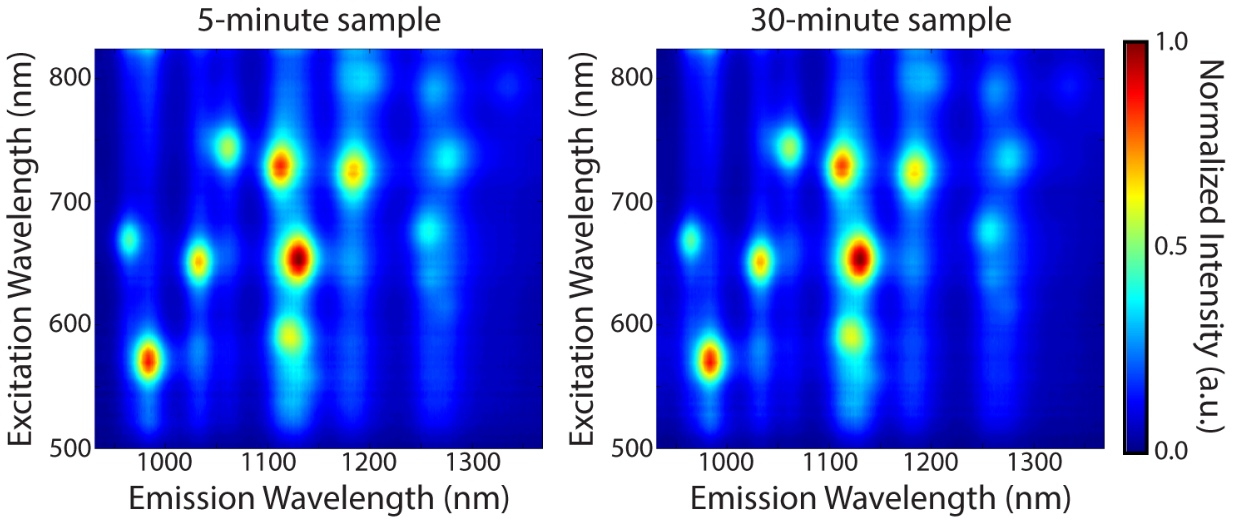


Figure S5. Two-dimensional photoluminescence excitation emission plots of the 5-minute sample and the 30-minute sample. Intensity was independently normalized for each plot.

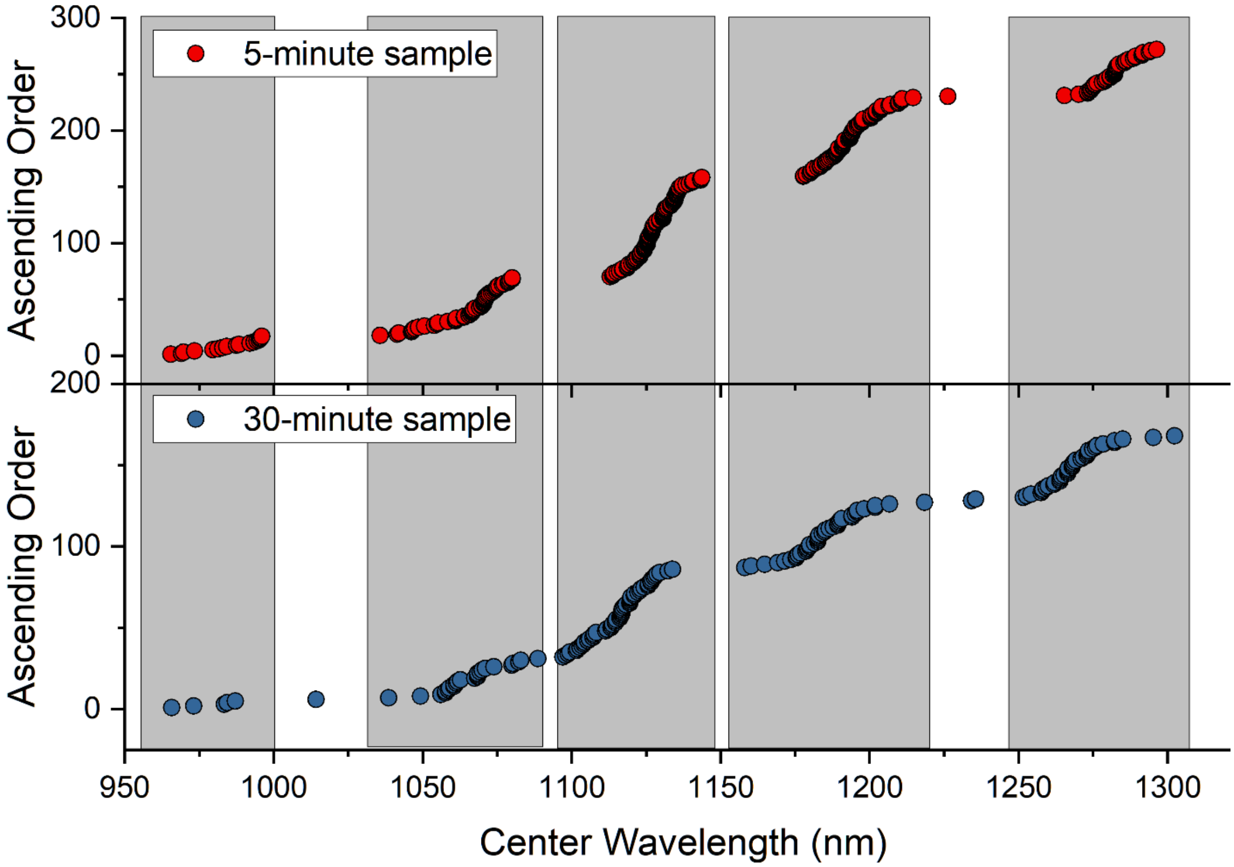


Figure S6. Clustering of emission center wavelength of individual surface-adsorbed ROIs from both samples. The shaded boxes indicate five emission bands. N = 272 for the 5-minute sample and N = 170 for the 30-minute sample.

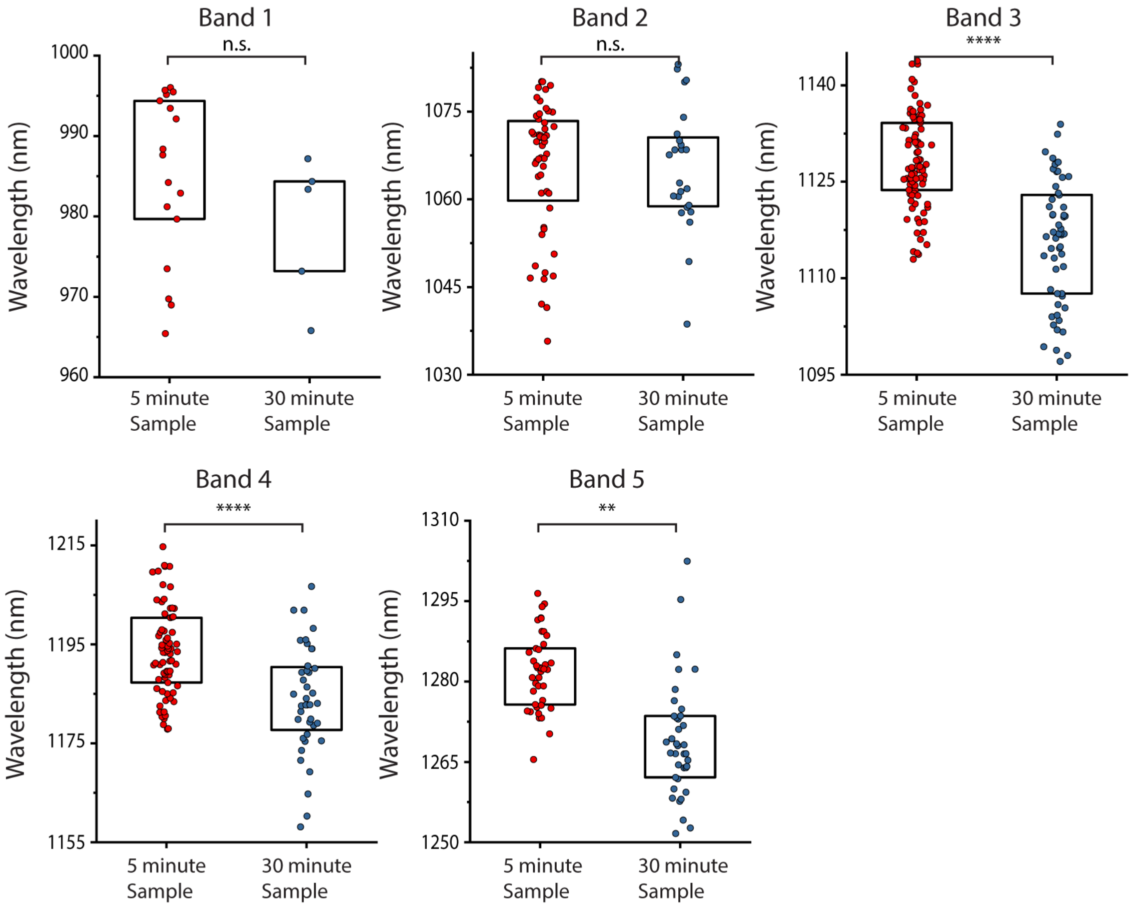


Figure S7. Emission center wavelength of surface-adsorbed SDC-SWCNT complexes in each band from both samples. Scatter plot of emission center wavelength of all individual ROIs emitting in each band from the two preparations. Boxes represent 25-75% of the data. Statistical comparisons are unpaired t-tests with Welch's correction. ** indicates < 0.01, **** indicates $p < 0.0001$.

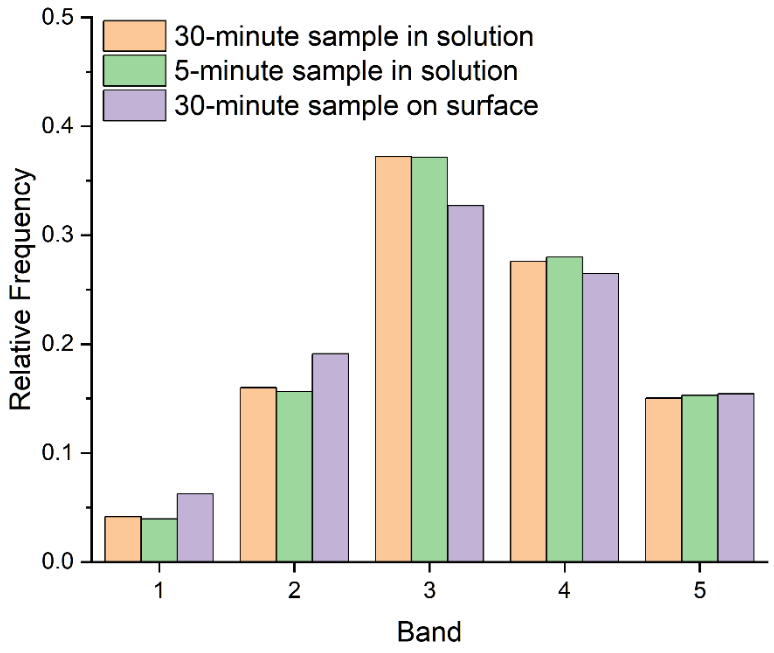


Figure S8. Population distributions of 5-minute and 30-minute samples. Solution population distribution quantified using the intensity of emission peaks corresponding to each band in the spectra obtained via hyperspectral microscopy of surface-adsorbed SWCNTs, as detailed in Fig. 2b.

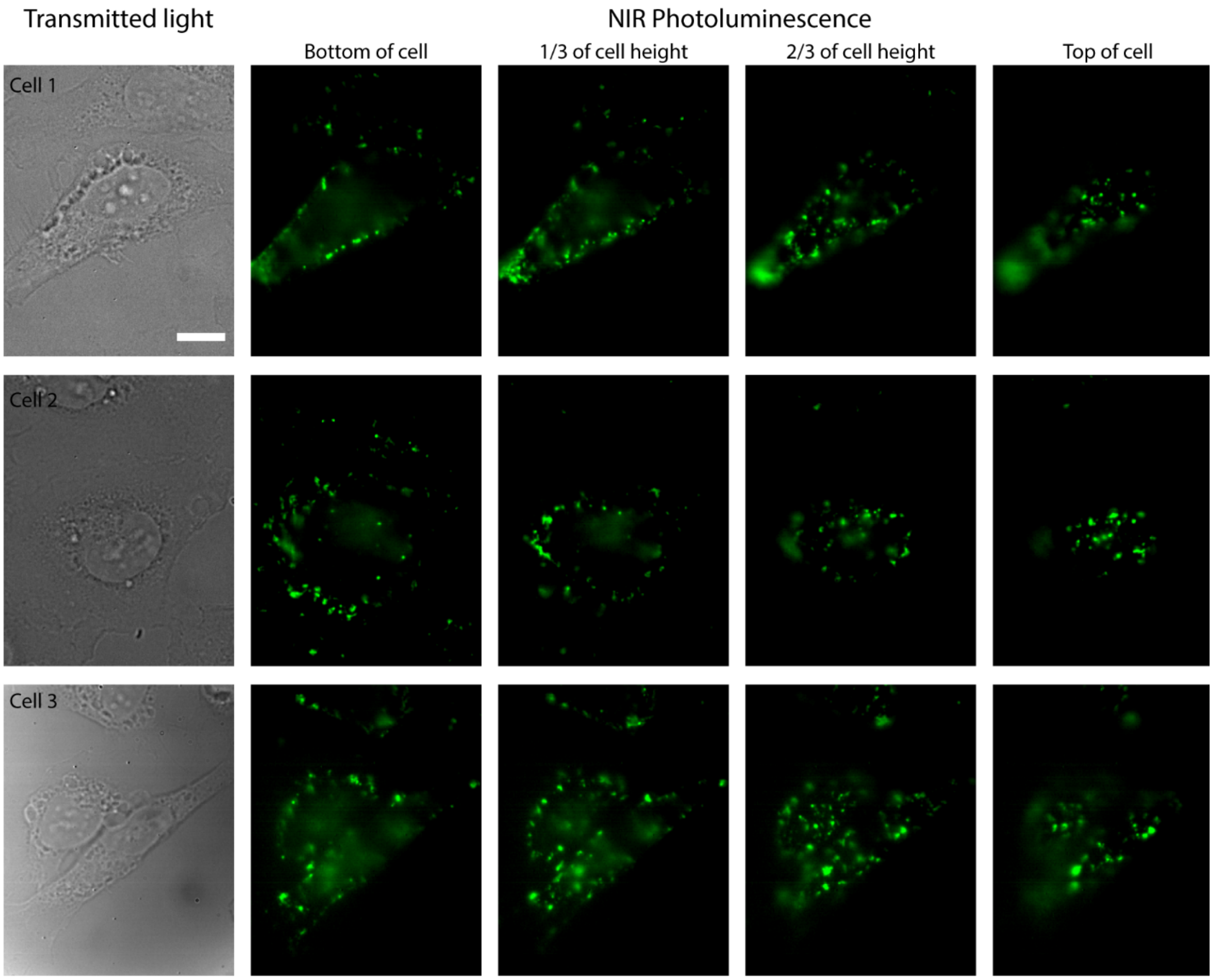


Figure S9. Photoluminescence imaging study of SDC-SWCNT distribution in HeLa cells. Transmitted light and NIR broadband images focusing on the bottom, 1/3$^{rd}$, 2/3$^{rd}$ and top of 3 typical HeLa cells incubated with 1 mg/L SDC-SWCNTs for 30 minutes. Scale bar is 10 µm.

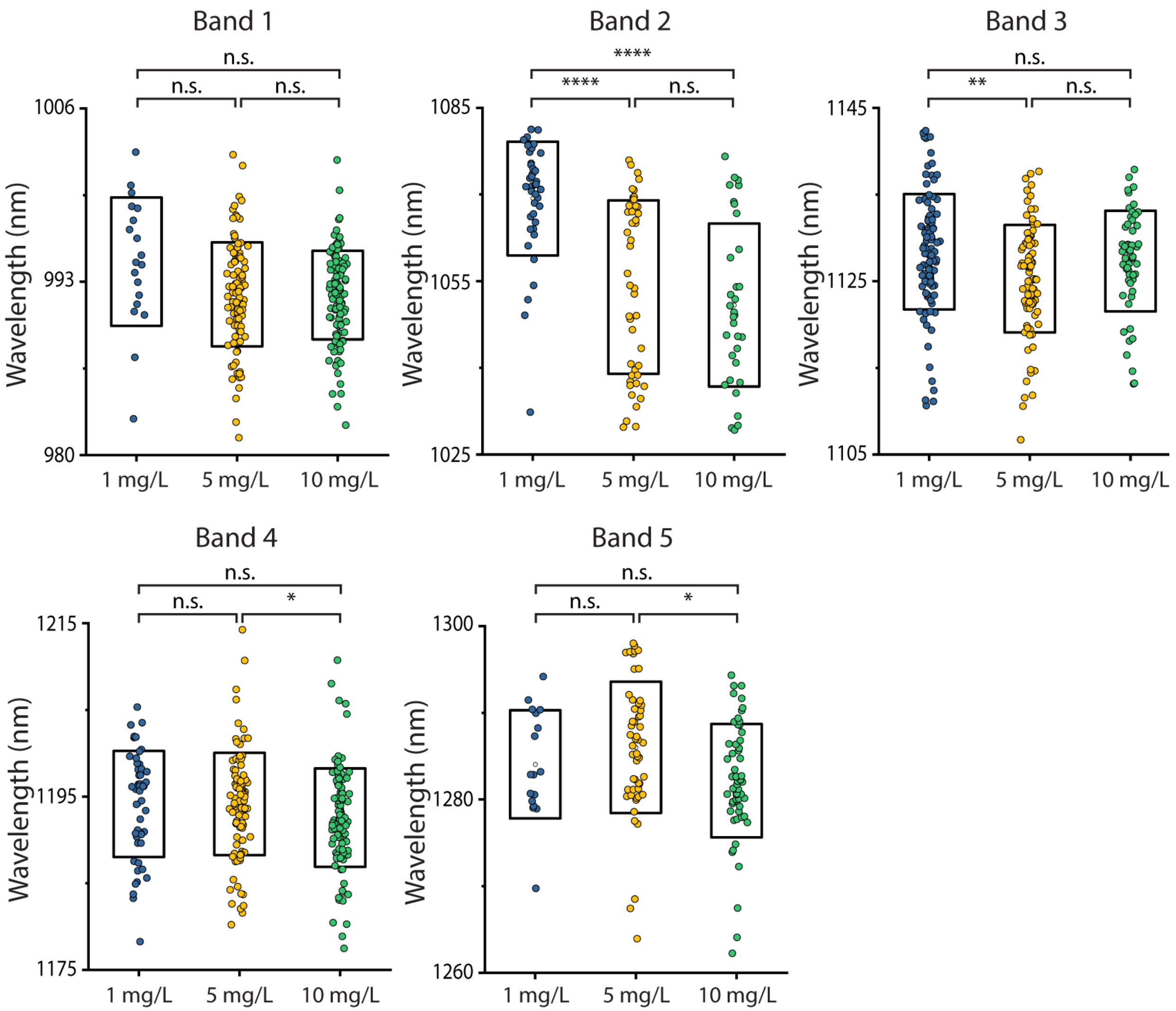


Figure S10. Emission wavelengths of SDC-SWCNT complexes in HeLa cells at 3 SDC-SWCNT loading concentrations. Center wavelength of individual ROIs in bands 1 to 5 (corresponding to the emission ranges defined in Table S4) for HeLa cells incubated with 1, 5 and 10 mg/L of SDC-SWCNTs for 30 minutes. One-way ANOVA was performed using Holm-Sidak's multiple comparison test. * = $p < 0.5$, ** = $p < 0.01$, **** = $p < 0.001$.

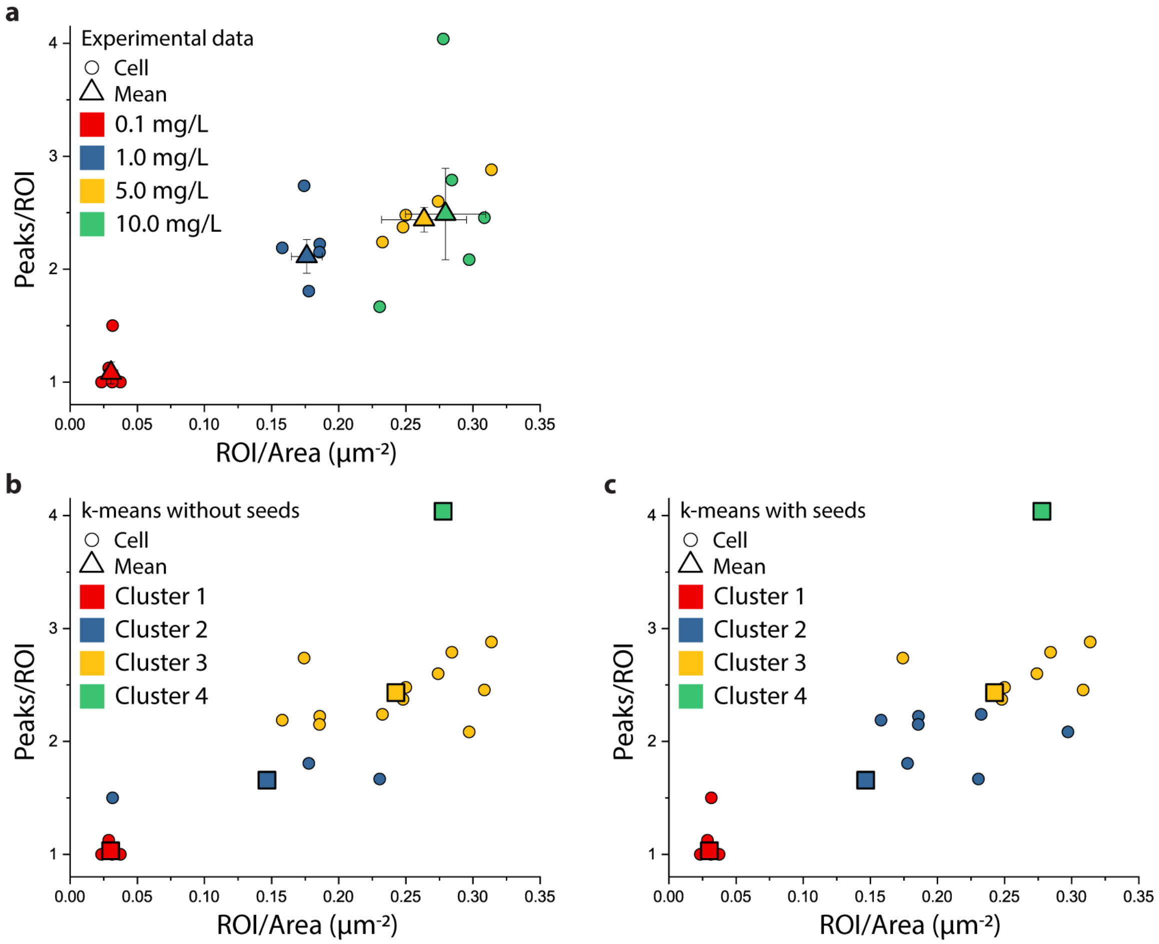


Figure S11. K-means analysis of concentration-dependent peaks per ROI vs. ROIs per μm$^2$ for single cells. (a) Experimental values of peaks per ROI vs ROI per area for cells incubated with 0.1, 1, 5 and 10 mg/L SDC-SWNT for 30-minutes (stock solution for cellular assays). (b) Clusters identified from the same set of data as in panel a using an unbiased k-means algorithm. (c) Clusters identified from the same data set as in panel a by a k - means algorithm using mean positions from the references data in panel a as starting seed locations.

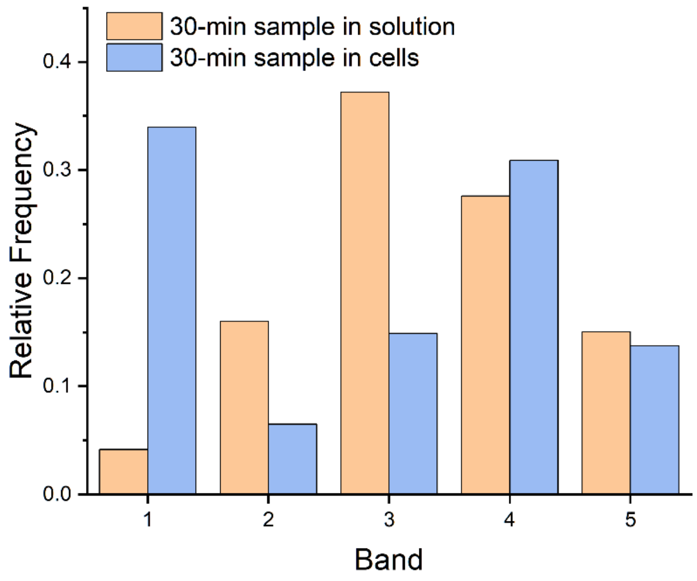


Figure S12. Population distribution of SDC-SWCNTs in solution and in HeLa cells. Solution population distribution of the 30-minute sample was measured using the intensity of emission peaks corresponding to each band in the emission spectra. The distribution in cells was obtained via hyperspectral microscopy of HeLa cells incubated with the 30-minute SDC-SWCNT sample (stock solution for cellular assays).

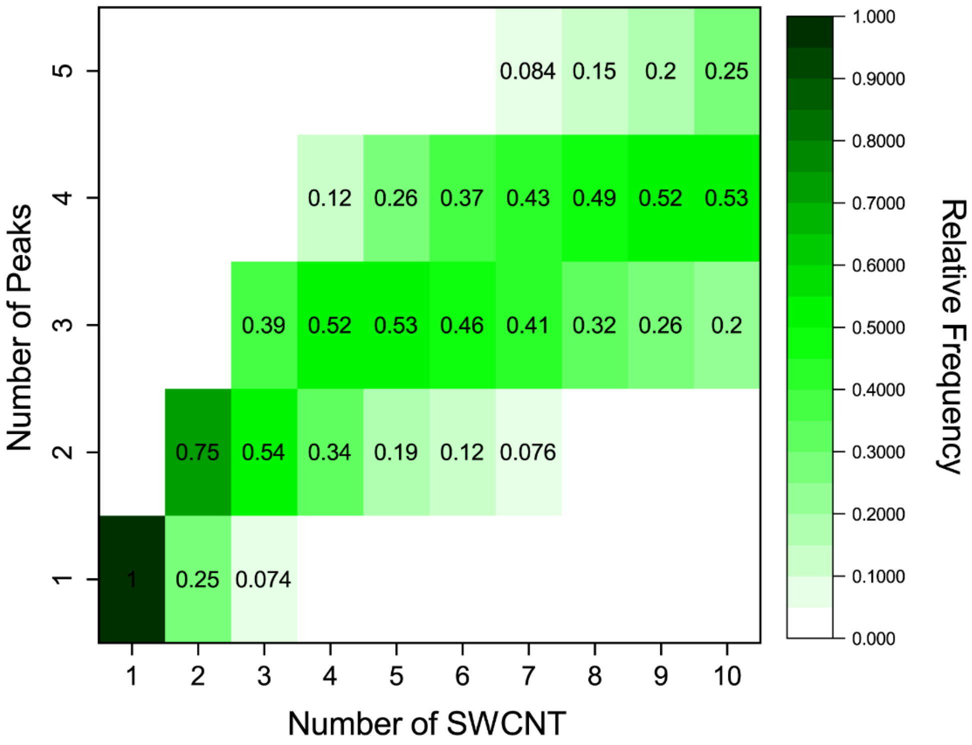


Figure S13. Heat map of the number of SWCNTs per ROI and the probability distribution of the number of emission peaks detected per ROI. This mapping relation is for HeLa cells incubated with the 30-minute SDC-SWCNT sample (stock solution for cellular assays), with photoluminescence emission acquired under 730 nm excitation. Values below 0.05 are not shown, for clarity.

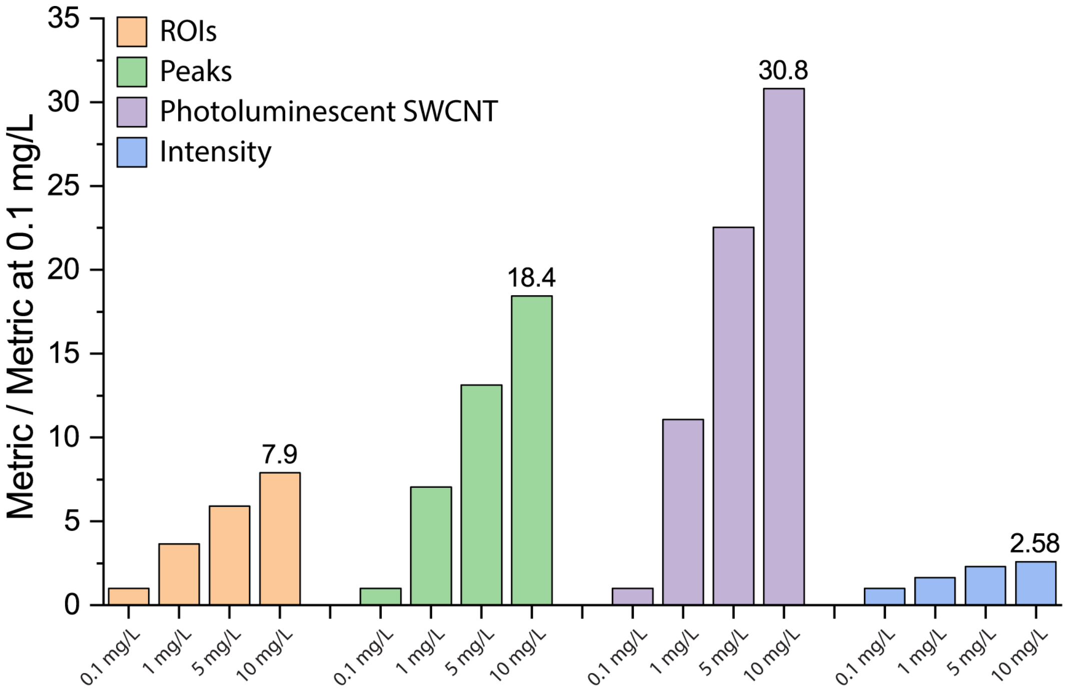


Figure S14. Comparison of four different metrics to quantify SWCNTs in cells. Each measurand is divided by its value at 0.1 mg/L SDC-SWCNT concentration to in order to compare the particle number counts (ROIs per cell, peaks per cell, photoluminescent SWCNTs per cell) with intensity. Photoluminescent intensity was calculated as the mean emission intensity over the area of each cell.

**Table S1: Optical Parameters from Photoluminescence Excitation Emission (PLE) Plots**

PLE Analysis - Normalized Intensity

| Chirality | Intensity – 5 min (a.u.) | Intensity – 30 min (a.u.) | % Difference (a.u) |
|---|---|---|---|
| (8,3) | 0.066423 ± 0.000102 | 0.068873 ± 0.000545 | ****, 3.7% increase |
| (6,5) | 0.12217 ± 0.000139 | 0.12445 ± 0.000568 | ****, 1.8% increase |
| (7,5) | 0.10175 ± 0.000113 | 0.10447 ± 0.000273 | ****, 2.6% increase |
| (10,2) | 0.078863 ± 0.000168 | 0.08011 ± 0.000411 | **, 1.6% increase |
| (9,4) | 0.12188 ± 0.000224 | 0.12109 ± 0.00049 | ns |
| (8,4) | 0.090087 ± 0.000182 | 0.089973 ± 0.000152 | ns |
| (7,6) | 0.14597 ± 0.0000586 | 0.14799 ± 0.000107 | ****, 1.4% increase |
| (8,6) | 0.09706 ± 0.000201 | 0.094167 ± 0.000298 | ****, 2.9% decrease |
| (9,5) | 0.056377 ± 0.000258 | 0.054717 ± 0.00026 | ***, 2.9% decrease |
| (10,5) | 0.042943 ± 0.000219 | 0.040873 ± 0.00026 | ****, 4.8% decrease |
| (8,7) | 0.05368 ± 0.000111 | 0.051987 ± 0.000166 | ***, 3.2% decrease |
| (9,7) | 0.022777 ± 0.0000524 | 0.021307 ± 0.000142 | **, 6.5% decrease |

PLE Analyses - Excitation Maximum

| Chirality | Excitation – 5 min (nm) | Excitation – 30 min (nm) | Difference (nm) |
|---|---|---|---|
| (8,3) | 668.47 ± 0.0248 | 668.21 ± 0.179 | *, 0.25 nm blue-shift |
| (6,5) | 570.66 ± 0.0323 | 570.62 ± 0.0328 | ns |
| (7,5) | 650.2 ± 0.026 | 650.16 ± 0.0442 | ns |
| (10,2) | 743.02 ± 0.0221 | 742.95 ± 0.00509 | ns |
| (9,4) | 728.09 ± 0.0146 | 728.08 ± 0.0492 | ns |
| (8,4) | 590.79 ± 0.0435 | 590.74 ± 0.044 | ns |
| (7,6) | 652.97 ± 0.0246 | 652.99 ± 0.00959 | ns |
| (8,6) | 723.82 ± 0.0237 | 723.72 ± 0.0176 | ns |
| (9,5) | 675.35 ± 0.0126 | 675.24 ± 0.0225 | ns |
| (10,5) | 789.37 ± 0.067 | 789.01 ± 0.051 | ***, 0.36 nm blue-shift |
| (8,7) | 734.19 ± 0.15 | 734.13 ± 0.0282 | ns |
| (9,7) | 793.66 ± 0.0398 | 793.23 ± 0.0786 | ****, 0.42 nm blue-shift |

PLE Analyses - Emission Maximum

| Chirality | Emission – 5 min (nm) | Emission – 30 min (nm) | Difference (nm) |
|---|---|---|---|
| (8,3) | 965.3 ± 0.00769 | 965.15 ± 0.0298 | ****, 0.15 nm blue-shift |
| (6,5) | 983.54 ± 0.02 | 983.4 ± 0.0366 | ****, 0.14 nm blue-shift |
| (7,5) | 1032.9 ± 0.00791 | 1032.8 ± 0.0058 | ****, 0.14 nm blue-shift |
| (10,2) | 1060.4 ± 0.0162 | 1060.5 ± 0.0152 | ****, 0.12 nm red-shift |
| (9,4) | 1113 ± 0.00321 | 1112.8 ± 0.016 | ****, 0.20 nm blue-shift |
| (8,4) | 1122 ± 0.00657 | 1121.8 ± 0.0116 | ****, 0.25 nm blue-shift |
| (7,6) | 1129.8 ± 0.00591 | 1129.6 ± 0.00953 | ****, 0.22 nm blue-shift |
| (8,6) | 1184.1 ± 0.00865 | 1183.8 ± 0.017 | ****, 0.34 nm blue-shift |
| (9,5) | 1258.5 ± 0.0123 | 1258.2 ± 0.0128 | ****, 0.28 nm blue-shift |
| (10,5) | 1264 ± 0.00662 | 1263.7 ± 0.0173 | ****, 0.37 nm blue-shift |
| (8,7) | 1275.5 ± 0.0157 | 1275.1 ± 0.0267 | ****, 0.36 nm blue-shift |
| (9,7) | 1335 ± 0.0213 | 1334.6 ± 0.0453 | ****, 0.41 nm blue-shift |

PLE Analyses – Emission Full-Width at Half-Maximum

| Chirality | FWHM – Emission (nm) 5 min | FWHM – Emission (nm) 30 min | Difference (nm) |
|---|---|---|---|
| (8,3) | 40.954 ± 0.0735 | 39.938 ± 0.623 | ns |
| (6,5) | 37.318 ± 0.0343 | 36.622 ± 0.167 | ns |
| (7,5) | 35.014 ± 0.0715 | 34.142 ± 0.154 | ns |
| (10,2) | 38.232 ± 0.229 | 36.985 ± 0.2 | ns |
| (9,4) | 39.683 ± 0.112 | 38.979 ± 0.0672 | ns |
| (8,4) | 75.518 ± 0.369 | 82.641 ± 2.3 | ****, 7.1 nm increase |
| (7,6) | 33.345 ± 0.0175 | 34.381 ± 0.127 | ns |
| (8,6) | 38.635 ± 0.0834 | 38.762 ± 0.139 | ns |
| (9,5) | 59.23 ± 1.05 | 48.148 ± 0.544 | ****, 11 nm decrease |
| (10,5) | 72.055 ± 2.14 | 71.697 ± 0.402 | ns |
| (8,7) | 28.379 ± 0.284 | 27.856 ± 0.266 | ns |
| (9,7) | 31.688 ± 1.82 | 36.802 ± 2.16 | **, 5.1 nm increase |

PLE Analyses – Excitation Full-Width at Half-Maximum

| Chirality | FWHM – Excitation (nm) 5 min | FWHM – Excitation (nm) 30 min | Difference (nm) |
|---|---|---|---|
| (8,3) | 102.54 ± 0.554 | 109.09 ± 0.838 | ns |
| (6,5) | 204.09 ± 8.17 | 213.07 ± 4.37 | ns |
| (7,5) | 126.88 ± 1.38 | 125.74 ± 0.799 | ns |
| (10,2) | 89.575 ± 0.751 | 85.981 ± 0.547 | ns |
| (9,4) | 95.657 ± 0.247 | 92.415 ± 0.416 | ns |
| (8,4) | 130.77 ± 1.38 | 131.65 ± 2.18 | ns |
| (7,6) | 160.24 ± 1.39 | 158.31 ± 1.64 | ns |
| (8,6) | 136.16 ± 1.79 | 135.89 ± 0.615 | ns |
| (9,5) | 122.95 ± 3.66 | 137.41 ± 6.43 | ns |
| (10,5) | 96.735 ± 5.37 | 106.07 ± 0.663 | ns |
| (8,7) | 52.348 ± 35.9 | 56.93 ± 0.0686 | ns |
| (9,7) | 64.274 ± 0.838 | 67.949 ± 4.11 | ns |

Table S2: Band edges and widths from k-means clustering of emission bands for SDC-SWCNT adsorbed on a surface

| Band | Starting Wavelength (nm) | Ending Wavelength (nm) | Band Size (nm) |
|---|---|---|---|
| 1 | 956 | 1020 | 64 |
| 2 | 1024 | 1092 | 68 |
| 3 | 1104 | 1160 | 56 |
| 4 | 1172 | 1232 | 60 |
| 5 | 1260 | 1312 | 52 |

Table S3: Optical parameters of SDC-SWCNT bands on surface

| Band | Wavelength (nm) – 5-minute sample | | Wavelength (nm) – 30-minute sample | |
|---|---|---|---|---|
| | Mean ± SEM | CI [25,75] | Mean ± SEM | CI [25,75] |
| 1 | 984.9 ± 2.519 | [979.7, 994.4] | 978.8 ± 4.014 | [973.2, 984.4] |
| 2 | 1065 ± 1.575 | [1060, 1073] | 1065 ± 2.151 | [1059, 1071] |
| 3 | 1128 ± 0.7657 | [1124, 1134] | 1116 ± 1.287 | [1108, 1123] |
| 4 | 1194 ± 1.073 | [1187, 1200] | 1184 ± 1.723 | [1178, 1190] |
| 5 | 1282 ± 1.079 | [1276, 1286] | 1269 ± 1.737 | [1262, 1274] |

| Band | Intensity (a.u.) – 5-minute sample | | Intensity (a.u.) – 30-minute sample | |
|---|---|---|---|---|
| | Mean ± SEM | CI [25,75] | Mean ± SEM | CI [25,75] |
| 1 | 2247 ± 234.9 | [1535, 3032] | 2188 ± 473.5 | [1505, 3011] |
| 2 | 2388 ± 185.3 | [1310, 3032] | 2021 ± 182.6 | [1331, 2583] |
| 3 | 2688 ± 205.3 | [1381, 3301] | 2440 ± 200.7 | [1501, 2982] |
| 4 | 2420 ± 224.2 | [1241, 2875] | 1947 ± 156.3 | [1267, 2326] |
| 5 | 2605 ± 228.3 | [1835, 3129] | 2410 ± 254.8 | [1395, 2955] |

**Supplementary Text 1: Details of computational model**

We developed a three-step computational method to approximate the number of emissive SWCNTs present in an ROI from the number of emission peaks: In the first step, a distribution of the nanotube population across the emission bands was experimentally determined via optical measurements. In the second step, a probabilistic relationship between the number of emissive peaks detected from an ROI and the number of SWCNT present in the ROI was estimated. In the final step, the experimentally determined distribution of the number of emission peaks was used to solve a system of linear equations for the number of emissive SWCNTs.

**Determining band distribution of the nanotube population**

Our model requires an accurate estimation of the experimentally-determined relative sizes of each emission band, under the same experimental conditions under which the hyperspectral data will be fit. In our experiments, we used a 730 nm laser to excite the sample. For solution measurements, we found the most accurate metric to be the fitted peak height of each emission band. For single-nanotube microscopy experiments, the number of peaks observed per emission band was a directly measure of the relative size of each emission band.

For SDC-SWCNT adsorbed on a surface, the relative sizes of the bands were:
[B1; B2; B3; B4; B5] = [0.0625; 0.1912; 0.3272; 0.3647; 0.1544]

**Monte Carlo simulation to map emission peaks to emissive SWCNT**

The process of distributing N emissive nanotubes into 5 emission bands of different sizes was simulated 5,000 times to estimate the resulting distribution of emissive nanotubes per band. Outputs from the first 5 runs are shown below for N = 5:

| Run | Band 1 | Band 2 | Band 3 | Band 4 | Band 5 |
|---|---|---|---|---|---|
| 1 | 0 | 3 | 0 | 1 | 1 |
| 2 | 1 | 0 | 2 | 1 | 1 |
| 3 | 0 | 2 | 1 | 1 | 1 |
| 4 | 1 | 1 | 3 | 0 | 0 |
| 5 | 0 | 1 | 1 | 1 | 2 |

This output was transformed according to the rules of peak reduction i.e., 1 or more emissive nanotubes per band result in 1 detected emission peak. The above table transforms as:

| Run | Band 1 | Band 2 | Band 3 | Band 4 | Band 5 |
|---|---|---|---|---|---|
| 1 | 0 → 0 | 3 → 1 | 0 → 0 | 1 → 1 | 1 → 1 |
| 2 | 1 → 1 | 0 → 0 | 2 → 1 | 1 → 1 | 1 → 1 |
| 3 | 0 → 0 | 2 → 1 | 1 → 1 | 1 → 1 | 1 → 1 |
| 4 | 1 → 1 | 1 → 1 | 3 → 1 | 0 → 0 | 0 → 0 |
| 5 | 0 → 0 | 1 → 1 | 1 → 1 | 1 → 1 | 2 → 1 |

For the 5 simulations above, summing each row provides the number of emissive SWCNT per ROI for the first table, and the number of emission peaks per ROI for the second table

| Run | Emissive SWCNT | Emission Peaks |
|---|---|---|
| 1 | 5 | 3 |
| 2 | 5 | 4 |
| 3 | 5 | 4 |
| 4 | 5 | 3 |
| 5 | 5 | 4 |

After 5,000 such simulations, the resulting relative frequency histograms for the number of emissive SWCNT present and the number of emission peaks detected are as follows:

| Number of SWCNT | Relative Frequency | Number of Peaks | Relative Frequency |
|---|---|---|---|
| 1 | 0 | 1 | 0.0040 |
| 2 | 0 | 2 | 0.1640 |
| 3 | 0 | 3 | 0.5330 |
| 4 | 0 | 4 | 0.2770 |
| 5 | 5 | 5 | 0.0220 |

For each value of N from 1 to 10, there is a corresponding probability distribution function for the number of emission peaks detected when N emissive SWCNT are present, for each ROI.

In terms of equation 5 from the main text,

$$N_5 \rightarrow 0.0040P_1 + 0.1640P_2 + 0.5330P_3 + 0.2770P_4 + 0.0220P_5$$

Where $P_1$, $P_2$, $P_3$, $P_4$ and $P_5$ are the probabilities of detecting 1,2,3,4 and 5 emission peaks when 5 emissive SWCNT are present in the ROI.

**Estimating distribution of emissive SWCNT from distribution of emission peaks**

The experimentally determined distribution of the number of emission peaks per ROI is used to estimate the underlying distribution of the number of emissive SWCNT.

For each N (from 1 to 10), there is a relationship of the form:

$$N_i \rightarrow b_1P_1 + b_2P_2 + b_3P_3 + b_4P_4 + b_5P_5$$

Where $i$ = 1 to 10.

We are solving for the opposite equation i.e., if P emission peaks are detected, what is the probability distribution of there being N emissive SWCNT present:

$$P_j \rightarrow a_1N_1 + a_2N_2 + a_3N_3 + a_4N_4 + a_5N_5 + a_6N_6 + a_7N_7 + a_8N_8 + a_9N_9 + a_{10}N_{10}$$

Where $j$ = 1 to 5.

The system of equations above are a form of constrained linear least squares problem. For a given set of $b_i$ (relative frequency histograms for the number of emission peaks detected), we solve the system to obtain a set of solutions $a_j$. To test the goodness of these fits, we used $a_j$ as the input to estimate the $b_i$ coefficients.

For the case of the 5-minute sample on the surface,

The input data is:

| **Input** | | **Simulation** | | **Goodness of Fit Check** | |
|---|---|---|---|---|---|
| Number of Peaks | Relative Frequency | Number of SWCNT | Relative Frequency | Number of Peaks | Relative Frequency |
| 1 | 0.693 | 1 | 0.63829 | 1 | 0.69298 |
| 2 | 0.212 | 2 | 0.21047 | 2 | 0.21217 |
| 3 | 0.075 | 3 | 0.04986 | 3 | 0.07447 |
| 4 | 0.018 | 4 | 0.07826 | 4 | 0.01804 |
| 5 | 0.002 | 5 | 0.00846 | 5 | 0.00224 |
| | | 6 | 0.00486 | | |
| | | 7 | 0.00341 | | |
| | | 8 | 0.00235 | | |
| | | 9 | 0.00215 | | |
| | | 10 | 0.0018 | | |

**Supplementary Text 2: Detailed calculation of nanotube concentration within single cells and single endosomes**

**Calculating loading SWCNT concentration from absorbance measurement**

Extinction coefficient $\varepsilon_{mass}$ = 0.02554 L/(mg cm)
Extinction coefficient $\varepsilon_{moles}$ = 2 x $10^9$ mL/(mol cm)

From the Beer-Lambert Law, A = ε x ι x c

For the same concentration of SDC-SWCNT solution,

0.02554 L/(mg cm) x 1 cm x $C_{mg}$ = 2 x $10^6$ L/(mol cm) x 1 cm x $C_{mol}$

1 mole SWCNT = 2 x $10^6$/(0.02554) mg = 78.3 x $10^6$ mg SWCNT

Therefore, 10 mg SWCNT = 0.13 μmoles
10 mg/L SWCNT = 130 nM

Loading concentration of SWCNT in molarity:
10 mg/L → 130 nM
5.0 mg/L → 65 nM
1.0 mg/L → 13 nM
0.1 mg/L → 13 nM

**Calculation of SWCNT concentration in a cell**

HeLa Cell Volume = 3,000 $\mu m^3$
At 10 mg/L, approximately 3,000 SWCNT per cell is the limiting value

Concentration of SWCNT = molarity = moles / volume = 3000 SWCNT per cell

1 mole = 6 x $10^{23}$ molecules → 3000 molecules = 3000 moles / 6 x $10^{23}$

Molarity = 3000 moles / (6 x $10^{23}$ x 3,000 $\mu m^3$) = 0.17 x $10^{-23}$ moles/$\mu m^3$ = 1.7 nM

Therefore at 10 mg/L loading concentration, 3000 SWCNT inside a cell = 1.7 nM

**Calculation of SWCNT concentration in an endosome**

HeLa endosome volume = 0.021 $\mu m^3$
At 10 mg/L, approximately 4 SWCNT per endosome is the limiting value

Concentration of SWCNT = molarity = moles / volume = 4 SWCNT per endosome

Molarity of SWCNT = 4 moles / (6 x $10^{23}$ x 0.021 $\mu m^3$) = 31.0 x $10^{-14}$ moles/$\mu m^3$ = 319 nM
